# Ensemble cryo-EM structures demonstrate human IMPDH2 filament assembly tunes allosteric regulation

**DOI:** 10.1101/798322

**Authors:** Matthew C. Johnson, Justin M. Kollman

## Abstract

Inosine monophosphate dehydrogenase (IMPDH) mediates the first committed step in guanine nucleotide biosynthesis and plays important roles in cellular proliferation and the immune response. The enzyme is heavily regulated to maintain balance between guanine and adenine nucleotide pools. IMPDH reversibly polymerizes in cells and tissues in response to changes in metabolic demand, providing an additional layer of regulatory control associated with increased flux through the guanine synthesis pathway. Here, we report a series of human IMPDH2 cryo-EM structures in active and inactive conformations, and show that the filament resists inhibition by guanine nucleotides. The structures define the mechanism of filament assembly, and reveal how assembly interactions tune the response to guanine inhibition. Filament-dependent allosteric regulation of IMPDH2 makes the enzyme less sensitive to feedback inhibition, explaining why assembly occurs under physiological conditions, like stem cell proliferation and T-cell activation, that require expansion of guanine nucleotide pools.

## Introduction

Ribonucleotides play a central role in cellular physiology, and complex regulatory networks maintain optimal nucleotide levels according to the variable metabolic state of the cell (Lane & Fan 2015). Under most conditions, cells rely on salvage pathways to regenerate degradation products and maintain nucleotide pools. However when nucleotide demand is high, for example during cellular proliferation, flux through de novo nucleotide biosynthesis pathways is up-regulated.

The universally conserved enzyme IMP dehydrogenase (IMPDH) catalyzes the first committed step in guanine nucleotide synthesis. Initiation of purine nucleotide biosynthesis is tightly regulated by downstream adenine and guanine nucleotide products. Balancing the flux through these parallel synthesis pathways, which share the precursor inosine monophosphate (IMP), is essential for cellular homeostasis (Fig 1A) (Allison & Eugui 2000). IMPDH is regulated transcriptionally, post-translationally, and allosterically (Hedstrom 2009). In vertebrates, two IMPDH isoforms, (83% identical in humans), have differential expression patterns (Collart & Huberman 1988; Natsumeda et al. 1990). IMPDH1 is constitutively expressed at low levels in most tissues, while IMPDH2 is generally upregulated in proliferating tissues (Senda & Natsumeda 1994; Jackson et al. 1975; Hager et al. 1995; Carr et al. 1993). In mice, knockout of IMPDH1 results in only very minor vision defects, whereas knockout of IMPDH2 is embryonic lethal (Gu et al. 2003; Aherne et al. 2004; Gu et al. 2000).

**Figure 1.**
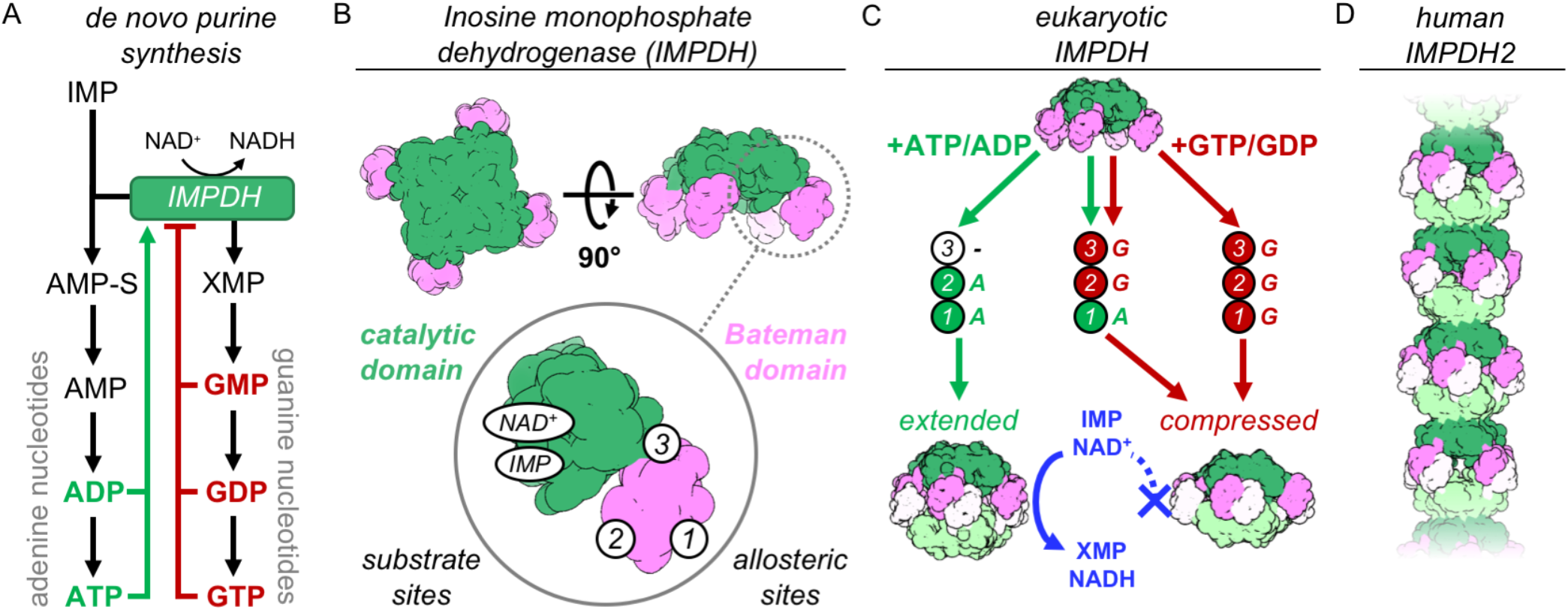
IMPDH structure and function. A) De novo purine nucleotide biosynthesis pathways. B) IMPDH consists of a catalytic domain with two substrate binding sites (green), and a regulatory Bateman domain with three allosteric binding sites on the Bateman domain numbered 1,2,3. C) Bound nucleotides promote regulatory domain dimerization, forming reversible IMPDH octamers that may active or inhibited. Opposing tetramers colored light green & light pink. D) Human IMPDH2 assembles into filaments composed of canonical octamers.

IMPDH reversibly assembles into filaments in vertebrate cells and tissues, providing an additional layer of regulation. Guanine deprivation induces assembly of IMPDH into micron-scale ultrastructures composed of bundled filaments, which disassemble once homeostasis is restored (Labesse et al. 2013; Calise et al. 2014; Thomas et al. 2012; Juda et al. 2014). Additional stimuli that alter IMPDH catalytic flux result in assembly, including increased intracellular IMP, and treatment with IMPDH inhibitors or other anti-proliferative drugs (Keppeke et al. 2018; Chang et al. 2015; Ji et al. 2006; Keppeke et al. 2015). In vivo, IMPDH assembly is seen mainly in cases of high nucleotide demand, for example assembly is correlated with proliferation of mouse induced pluripotent stem cells and human T-cell activation (Keppeke et al. 2018; Duong-Ly et al. 2018; Calise et al. 2018). However, the molecular mechanisms that underlie assembly-dependent regulation of IMPDH have not been described.

The catalytic mechanism and the structures of IMPDH monomers and defined oligomers are well-described (Hedstrom 2009). IMPDH forms stable tetramers, each protomer consisting of a catalytic domain and a regulatory Bateman domain (Fig 1B). The enzyme converts IMP to xanthosine monophosphate (XMP) and requires NAD^+^ as a cofactor for a complex multi-stage catalysis requiring rearrangements of active site loops that close around the substrate upon binding, and then must reopen to allow release of product. The regulatory domain contains three allosteric sites that bind adenine and guanine nucleotides. Sites 1 and 2 are canonical cystathionine beta synthase motifs that bind either ATP/ADP or GTP/GDP, and are conserved among IMPDH homologues (Scott et al. 2004; Ignoul & Eggermont 2005; Baykov et al. 2011; Ereño-Orbea et al. 2013). Site 3 is a non-canonical site located at the interface between domains that binds only GTP/GDP (Buey, Ledesma-Amaro, Velázquez-Campoy, et al. 2015).

In eukaryotes, adenine and guanine nucleotides allosterically modulate IMPDH activity by altering oligomeric state (Buey et al. 2017; Fernández-Justel et al. 2019). IMPDH tetramers reversibly assemble into octamers when nucleotides bind the two canonical sites and drive dimerization of Bateman domains (Fig 1C). Bound adenines promote an extended conformation in which the active sites are free to open and close as needed for catalysis. GTP/GDP binding induces a compressed conformation by changing the relative orientation of the two domains. This brings the active sites of opposing tetramers tightly together, forming an interdigitated pseudo beta-barrel that prevents reopening and product release and inhibits IMPDH activity by a mechanism described as a “conformational switch” between extended and compressed states (Buey, Ledesma-Amaro, Velázquez-Campoy, et al. 2015; Buey et al. 2017; Fernández-Justel et al. 2019). Because inhibition requires interactions between two opposing tetramers, this switch is functionally relevant in only the octameric state.

In vitro treatment with ATP or GTP induces assembly of human IMPDH into filaments composed of stacked octamers interacting through their catalytic domains (Fig. 1D) (Labesse et al. 2013; Anthony et al. 2017; Fernández-Justel et al. 2019). Filament segments can both extend and compress, and assembly does not have a direct effect on the activity of IMPDH2 (Anthony et al. 2017). Importantly, mutations that block assembly of IMPDH filaments in vitro also prevent assembly of the large IMPDH bundles observed in cells, supporting the functional relevance of in vitro reconstituted filaments.

In this study, we present a series of near-atomic resolution cryo-electron microscopy (cryo-EM) structures of human IMPDH2. Structures of the enzyme treated with multiple combinations of substrates (IMP, NAD+) and allosteric effectors (ATP, GTP), in both filament and non-filament assembly states, demonstrate the extreme conformational plasticity of the enzyme. These structures define the interactions that drive filament assembly, and we show that in vitro IMPDH2 filament assembly is sensitive to the same conditions that promote assembly in cells: high IMP levels and low guanine nucleotide levels (Keppeke et al. 2018). Finally, we show that filament assembly tunes sensitivity to GTP inhibition by stabilizing a conformation that reduces affinity for GTP.

## Results

### IMPDH2 filaments are conformationally heterogeneous

We first characterized IMPDH2 filaments assembled in vitro by addition of ATP (Anthony et al. 2017). The affinity of IMPDH2 for ATP has not been directly measured but we found ATP concentrations as low as 1 μM sufficient to induce assembly (Fig. 2A). We prepared cryo-EM grids of ATP-assembled filaments and found that in the absence of other ligands, IMPDH2 filaments are extremely flexible (Fig. S2A). Two-dimensional class averages confirmed our prior observation from negative stain that the filaments are composed of conformationally heterogenous octamers stacked head-to-head, resulting in flexible filaments with variable rise and radius of curvature (Fig. 2B). Because these deviations from ideal helical symmetry severely limited attempts at image processing by traditional iterative helical real-space reconstruction, we attempted to produce more structurally homogeneous filaments by addition of IMP or NAD^+^, which stabilize the flexible active site loops (Sintchak et al. 1996). As previously reported, substrates did not have a direct effect on IMPDH2 filament assembly, and filament assembly did not directly affect enzymatic activity (Figs. 2C-D). Unfortunately, addition of IMP and NAD^+^, either alone or in combination, did not significantly reduce filament flexibility (Fig. S1B-D). Two-dimensional class averages of helical segments again exhibited varying degrees of curvature and showed no correlation of structural states between neighboring IMPDH2 octamers, preventing successful three-dimensional processing with conventional helical approaches (Figs 2E-G).

**Figure 2.**
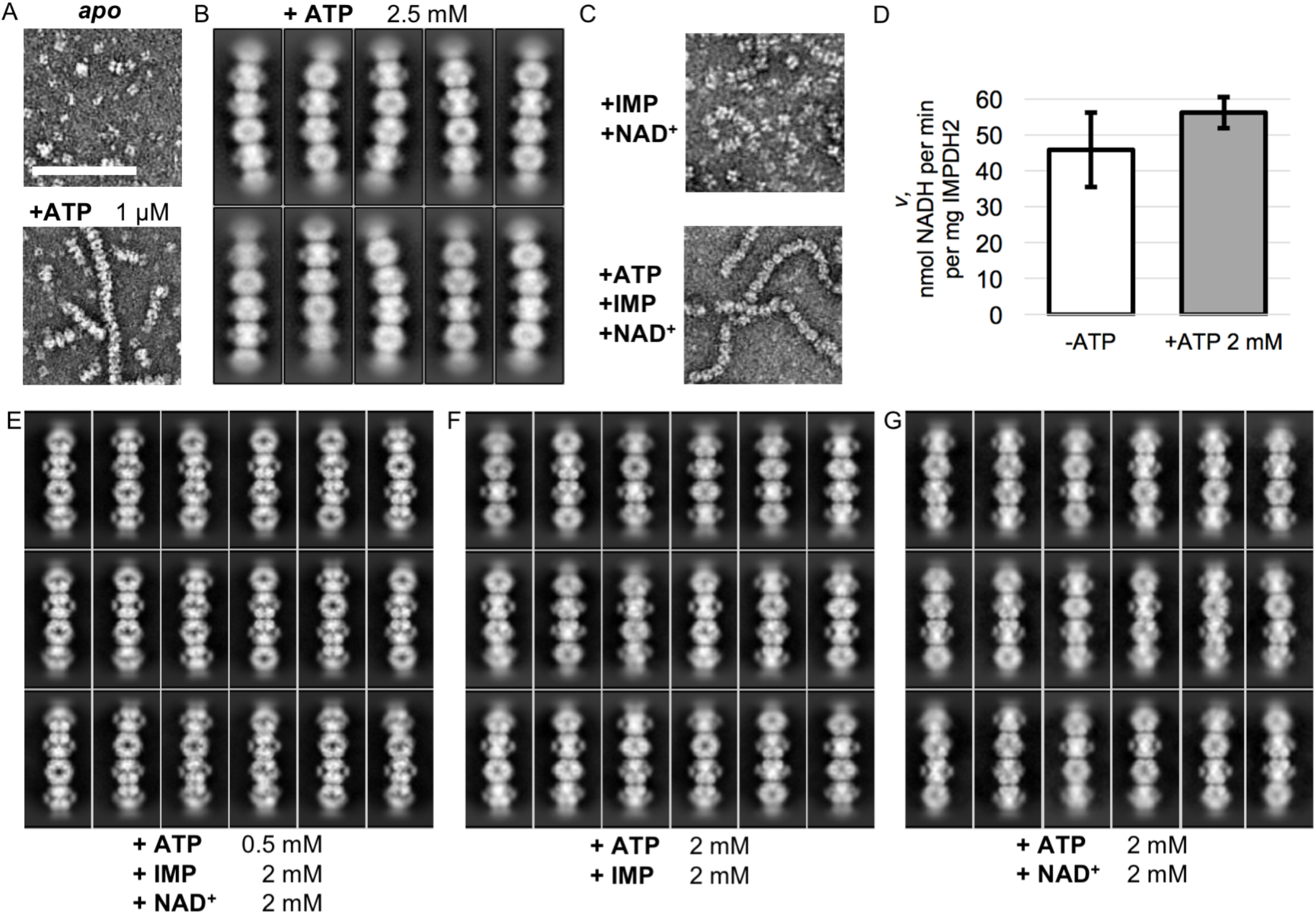
Electron microscopy of uninhibited IMPDH2 filaments. A) Negative stain EM of purified human IMPDH2. Treatment with 1 μM ATP induces filament assembly. Scale bar 100 nm. B) Representative 2D class averages from the +ATP cryo-EM dataset. C) Negative stain EM of actively catalyzing IMPDH2 (both substrates present at 2 mM), with and without 2 mM ATP. D) Initial velocity of enzyme with and without ATP (2 mM of both substrates). Average of three replicates, error bars +/-1 S.D. E-G) Representative 2D class averages from the three uninhibited enzyme cryo-EM datasets.

From these 2D class averages we observed that filament flexibility was due to variations in the conformation of filament segments, but that the interface between segments did not vary, and as a consequence this region was better resolved (Fig. 3A). Focused refinement of the filament assembly interface region alone provided a valuable foothold in resolving high resolution structures of all regions of the filaments. We developed a workflow for a single-particle style approach to reconstruction of inherently heterogeneous helical protomers, combining density subtraction, symmetry expansion, focused classification and focused refinement to isolate, classify, and reconstruct discrete sections of filaments for structural characterization (Fig. S1E-G). This approach provided multiple structures for two different focused regions of IMPDH2 filaments: full octameric segments, and the paired catalytic domain tetramers that constitute the filament assembly interface (Fig. 3B).

**Figure 3.**
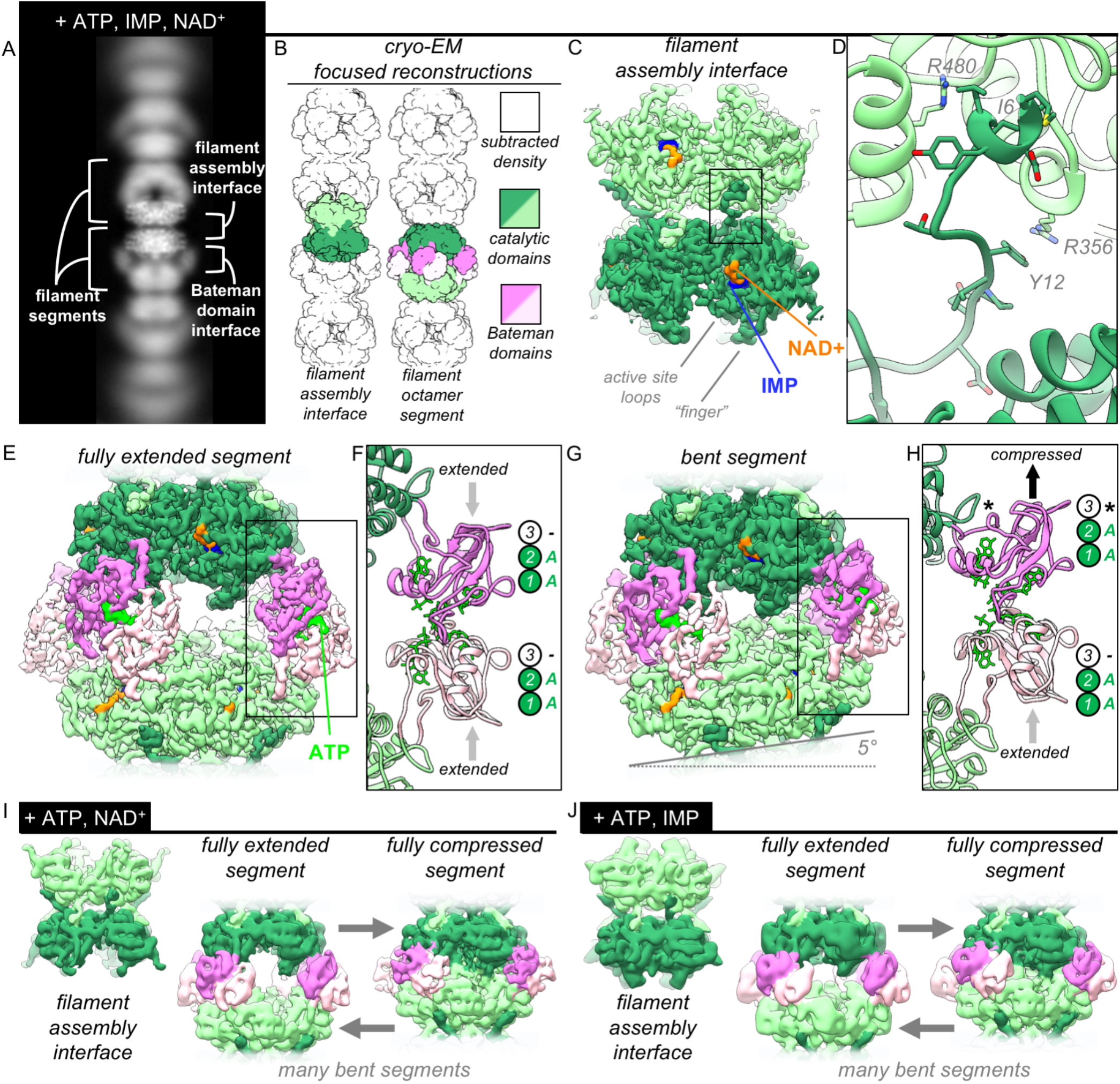
The structures of uninhibited IMPDH2 filaments. A) Cryo-EM of IMPDH2 filaments with both substrates (representative 2D class average). B) We resolved two types of structures from IMPDH filaments: the consensus filament assembly interface, and various conformations of filament segments. C) Cryo-EM density for the ATP/IMP/NAD+ consensus filament assembly interface, consisting of two tetramers bound back-to-back (dark and light green). D) The filament assembly interface is mediated by the vertebrate-specific N-terminus, in particular a key bridge between Y12 and R356. E) Cryo-EM density for the ATP/IMP/NAD+ fully extended filament segment. Opposing catalytic tetramers (dark and light green), are held separate by their symmetrically extended Bateman domains (dark and light pink). ATP (bright green) is resolvable in the Bateman domains. F) In the fully extended Bateman domains, sites 1 and 2 are occupied by ATP, and site 3 is unformed. G) Cryo-EM density for the best resolved ATP/IMP/NAD+ bent filament segment, in which the two catalytic tetramers are not parallel. H) Filament segment bending results from asymmetric compression of Bateman domains. In this reconstruction, one protomer from each of the two tetramers is compressed, and allosteric site 3 is formed, but unoccupied (black asterisk). I) Summary of the ATP/NAD+ cryo-EM dataset. The filament assembly interface is unchanged, and filament segments varied from fully extended, to bent, to fully compressed. In the absence of IMP, the flexible active site loops are disordered. J) Summary of the ATP/IMP cryo-EM dataset.

### The IMPDH2 filament assembly interface is well-ordered

To define the interactions that drive assembly of catalytically active IMPDH2 filaments, we obtained cryo-EM structures of IMPDH2 in three liganded states: 1) ATP/IMP/NAD^+^, 2) ATP/NAD^+^, and 3) ATP/IMP. The three datasets were qualitatively similar. However, because the structures refined to higher resolution when both substrates were present, we focus our analysis here on the ATP/IMP/NAD^+^ dataset. The complete image processing workflow applied to this dataset (Fig. S2A) was also applied to the two single-substrate datasets. From the ATP/IMP/NAD^+^ data we resolved many different structural classes, three of which reached high resolution: the filament assembly interface, and two different reconstructions of octameric filament segments (Table 1). We resolved many conformations of filament segments, however for all segments the filament assembly interface was identical. Therefore the filament assembly interface structure is a dataset consensus structure, an average of every segment included in the dataset. We did not observe any conformational cooperativity in the compressed/extended conformational equilibrium between IMPDH2 octamers sharing a filament assembly interface.

**Table 1.**
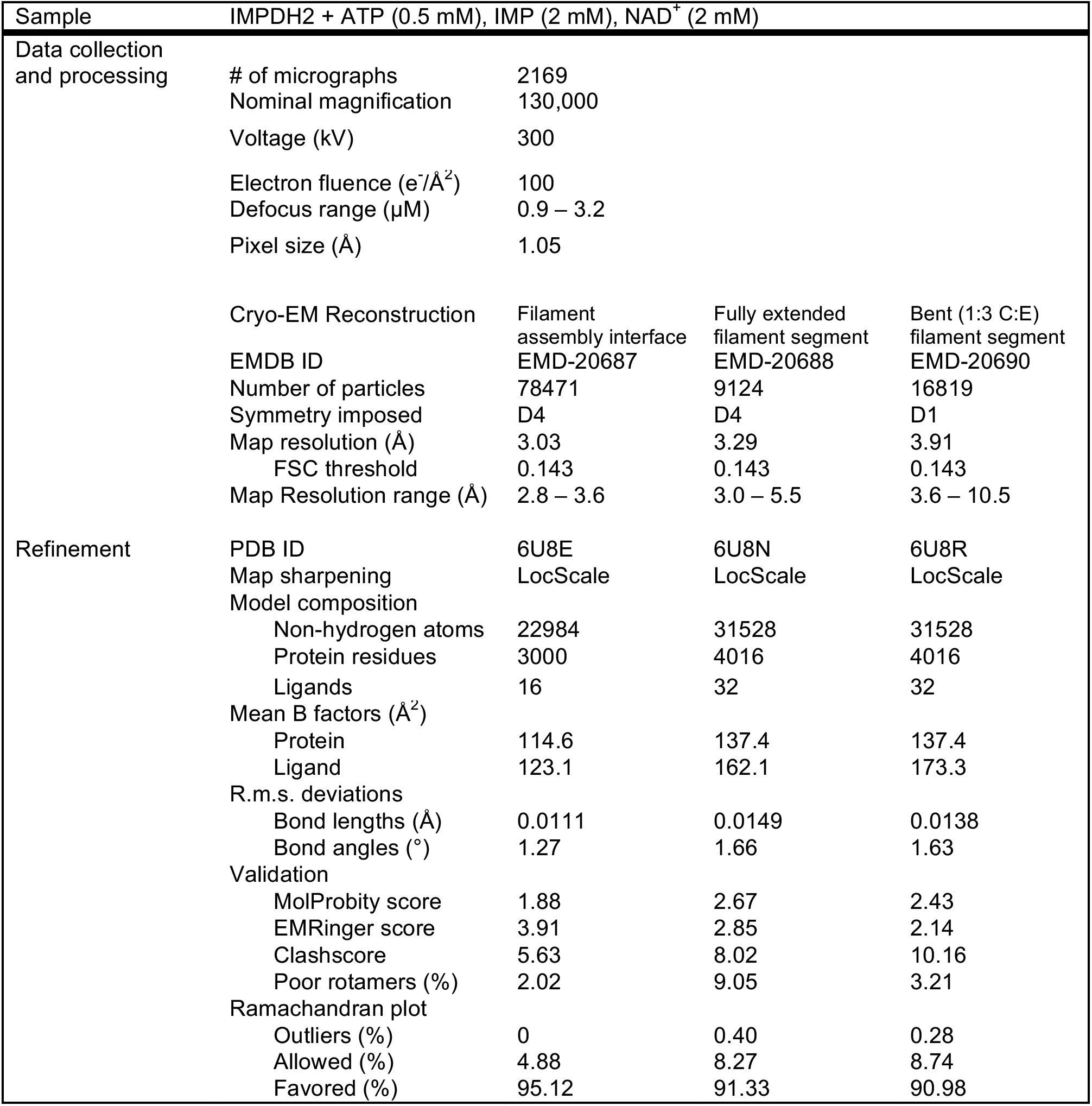
Statistics of cryo-EM data collection, reconstruction and model refinement for the ATP/IMP/NAD^+^ dataset.

| Sample | IMPDH2 + ATP (0.5 mM), IMP (2 mM), NAD<sup>+</sup> (2 mM) |  |  |  |
| --- | --- | --- | --- | --- |
| Data collection and processing | # of micrographs | 2169 |  |  |
|  | Nominal magnification | 130,000 |  |  |
|  | Voltage (kV) | 300 |  |  |
|  | Electron fluence (e<sup>-</sup>/Å<sup>2</sup>) | 100 |  |  |
|  | Defocus range (μM) | 0.9 – 3.2 |  |  |
|  | Pixel size (Å) | 1.05 |  |  |
|  | Cryo-EM Reconstruction | Filament assembly interface | Fully extended filament segment | Bent (1:3 C:E) filament segment |
|  | EMDB ID | EMD-20687 | EMD-20688 | EMD-20690 |
|  | Number of particles | 78471 | 9124 | 16819 |
|  | Symmetry imposed | D4 | D4 | D1 |
|  | Map resolution (Å) | 3.03 | 3.29 | 3.91 |
|  | FSC threshold | 0.143 | 0.143 | 0.143 |
|  | Map Resolution range (Å) | 2.8 – 3.6 | 3.0 – 5.5 | 3.6 – 10.5 |
| Refinement | PDB ID | 6U8E | 6U8N | 6U8R |
|  | Map sharpening | LocScale | LocScale | LocScale |
|  | Model composition |  |  |  |
|  | Non-hydrogen atoms | 22984 | 31528 | 31528 |
|  | Protein residues | 3000 | 4016 | 4016 |
|  | Ligands | 16 | 32 | 32 |
|  | Mean B factors (Å<sup>2</sup>) |  |  |  |
|  | Protein | 114.6 | 137.4 | 137.4 |
|  | Ligand | 123.1 | 162.1 | 173.3 |
|  | R.m.s. deviations |  |  |  |
|  | Bond lengths (Å) | 0.0111 | 0.0149 | 0.0138 |
|  | Bond angles (°) | 1.27 | 1.66 | 1.63 |
|  | Validation |  |  |  |
|  | MolProbity score | 1.88 | 2.67 | 2.43 |
|  | EMRinger score | 3.91 | 2.85 | 2.14 |
|  | Clashscore | 5.63 | 8.02 | 10.16 |
|  | Poor rotamers (%) | 2.02 | 9.05 | 3.21 |
|  | Ramachandran plot |  |  |  |
|  | Outliers (%) | 0 | 0.40 | 0.28 |
|  | Allowed (%) | 4.88 | 8.27 | 8.74 |
|  | Favored (%) | 95.12 | 91.33 | 90.98 |

We determined the structure of the ATP/IMP/NAD^+^ consensus filament assembly interface at 3.0 Å resolution (Figs. 3C, S2B-C). This D4 symmetric region is composed of two symmetrically opposed catalytic domain tetramers. The interface between tetramers is formed by the 12 amino-terminal residues of eight protomers, which each extend from the core of the molecule to bind one catalytic domain on the opposite tetramer, in a shallow surface groove formed by a short helix (476-485), two beta strands (51-63), and two short loops (355-360, 379-380) (Fig. 3D). A key tyrosine/arginine interaction (Y12/R356’) anchors the attachment; mutation of either of these residues to alanine was previously shown to abolish assembly, both in vivo and in vitro (Anthony et al. 2017). Residues 1-7 make mostly hydrophobic interactions with the catalytic domain, and an embedded arginine (R480’) is positioned to hydrogen bond to the I6 backbone carbonyl. These interactions are reciprocal - a protomer extends its N-terminus to a partner and receives the N-terminus of the same partner, creating four pairs of symmetrical interactions at the interface. The overall surface area buried by a single interaction is 2,300 Å^2^, with a total buried area of 9,200 Å^2^ for the multimeric interface.

The N-terminal tail of IMPDH, corresponding to residues 1-28 in IMPDH2, is the least conserved part of the protein, with large variation in length and sequence between phyla, as well as conformational variation among known structures (Fig. S3A) (Trapero et al. 2018; Makowska-Grzyska et al. 2015; Kim et al. 2017; Prosise & Luecke 2003; Osipiuk et al. 2014; Labesse et al. 2013; Buey, Ledesma-Amaro, Balsera, et al. 2015; Fernández-Justel et al. 2019). The residues involved in filament contacts, however, are conserved among chordates, consistent with the fact that IMPDH polymerization has only been reported in vertebrates (Fig. S3B). In previous crystal structures of human and fungal IMPDH, residues 1-11 are unresolved and residues 12-28 are well resolved. In these structures, Asp16 anchors the tail in place through ionic interactions with Arg341/Lys349 of a neighboring protomer, while Val13 and Pro14 make hydrophobic contacts. In the filament structure, however, the Val13/Pro14 contacts are broken and the entire tail is rotated about 30° to position Tyr12 to contact Arg356 across the filament interface (Fig. S3C).

In this consensus structure, the remainder of the catalytic domain is nearly identical to a structure of human hIMPDH2 bound to competitive inhibitors in a previously determined crystal structure (PDB 1nf7, backbone RMSD 0.641 Å for residues 18-107, 245-417, & 439-514) (Sintchak et al. 1996). The active site is well-resolved, except for one loop (residues 421-436). There is strong density in the both the IMP and NAD^+^ binding sites. We have modelled these ligands as IMP and NAD^+^, however because the enzyme filaments were actively turning over when flash frozen, we have likely captured a mixture of substrate-, intermediate-, and product-bound states. Attempts at focused classification of the active site to structurally isolate these states were unsuccessful.

### The extended conformation is an ensemble of flexible states

We resolved multiple conformations of filament segments in the ATP/IMP/NAD^+^ dataset (Fig. S2D). The most well-resolved was a D4-symmetric, fully extended octamer (3.3 Å, Figs. 3E, S2E-H). The eight Bateman domains are symmetrically extended with respect to their catalytic domains, giving the octameric segment a helical rise of 118 Å; a symmetric helix of these segments would have a twist of 36°. There is strong density at both of the canonical ATP-binding sites within the Bateman domain (Fig. 3F), consistent with previous crystal structures of ATP/ADP-bound fungal and bacterial IMPDH (Buey et al. 2017; Labesse et al. 2013).

We also resolved several bent filament segment structures, the best resolved of which reached 3.9 Å (Figs. 3G, S2I-L). This bent octamer contains two identical tetramers. For each, three protomers are in extended conformations, while one protomer is in the compressed conformation. Due to the symmetry of the octomer, the lone compressed protomer of each tetramer forms a Bateman domain dimer with an extended protomer of the opposing tetramer (Fig. 3H, Fig. S3D). From each of the lower resolution ATP/IMP and ATP/NAD^+^ datasets, we also observed a single consensus filament assembly interface and many different filament segment classes, including fully extended and bent segments, as well as fully compressed segments (Figs. 3I-J, Tables 2-3). To our knowledge, these are the first structure of IMPDH in the compressed conformation in the absence of guanine nucleotides. While the two canonical ATP sites are occupied, the third Bateman binding site (which forms only in the compressed state and is GTP/GDP-specific) remains unoccupied. This suggests that protomers within ATP-bound IMPDH filaments readily sample the compressed conformation, and that GTP binding to site 3 selectively stabilizes this state. Further, the ability of a compressed protomer to form a Bateman dimer with an extended partner demonstrates a lack of conformational cooperativity across the Bateman domain interface.

**Table 2.** Statistics of cryo-EM data collection, reconstruction and model refinement for the ATP/NAD^+^ dataset.

| Sample | IMPDH2 + ATP (2 mM), NAD<sup>+</sup> (2 mM) |  |  |  |
| --- | --- | --- | --- | --- |
| Data collection and processing | # of micrographs | 2289 |  |  |
|  | Nominal magnification | 130,000 |  |  |
|  | Voltage (kV) | 300 |  |  |
|  | Electron fluence (e<sup>-</sup>/Å<sup>2</sup>) | 100 |  |  |
|  | Defocus range (μM) | 0.5 – 7.0 |  |  |
|  | Pixel size (Å) | 1.062 |  |  |
|  | Cryo-EM Reconstruction | Filament assembly interface | Fully extended filament segment | Fully compressed filament segment |
|  | EMDB ID | EMD-20718 | EMD-20716 | EMD-20709 |
|  | Number of particles | 57979 | 13266 | 3394 |
|  | Symmetry imposed | D4 | D4 | D4 |
|  | Map resolution (Å) | 4.1 | 4.5 | 4.5 |
|  | FSC threshold | 0.143 | 0.143 | 0.143 |
|  | Map Resolution range (Å) | 3.8 – 7.8 | 4.3 – 7.0 | 4.2 – 7.3 |

**Table 3.** Statistics of cryo-EM data collection, reconstruction and model refinement for the ATP/IMP dataset.

| Sample | IMPDH2 + ATP (2 mM), IMP (3 mM) |  |  |  |
| --- | --- | --- | --- | --- |
| Data collection and processing | # of micrographs | 2178 |  |  |
|  | Nominal magnification | 105,000 |  |  |
|  | Voltage (kV) | 300 |  |  |
|  | Electron fluence (e<sup>-</sup>/Å<sup>2</sup>) | 100 |  |  |
|  | Defocus range (μM) | 0.6 – 4.2 |  |  |
|  | Pixel size (Å) | 1.366 |  |  |
|  | Cryo-EM Reconstruction | Filament assembly interface | Fully extended filament segment | Fully compressed filament segment |
|  | EMDB ID | EMD-20723 | EMD-20722 | EMD-20720 |
|  | Number of particles | 92587 | 9549 | 9648 |
|  | Symmetry imposed | D4 | D4 | D4 |
|  | Map resolution (Å) | 4.4 | 7.1 | 4.9 |
|  | FSC threshold | 0.143 | 0.143 | 0.143 |
|  | Map Resolution range (Å) | 4.2 – 6.7 | 5.5 – 8.1 | 4.2 – 10.5 |

### The balance of substrate and downstream product regulates filament assembly

We and others previously reported that GTP can stabilize compressed IMPDH2 filaments or drive their disassembly, depending on what other ligands are present (Buey et al. 2017; Anthony et al. 2017; Duong-Ly et al. 2018). To understand how ligand status of the active site tunes the response to GTP, we systematically explored the effects of different ligand combinations on filament assembly. In the absence of substrate, GTP induces disassembly of filaments, but as little as 10 μM IMP inhibits disassembly (Figs. 4A-B). Pre-treatment of filaments with IMP prevents disassembly by GTP, and addition of IMP promotes reassembly of filaments previously disassembled by GTP (Figs. S4A-B).

**Figure 4.**
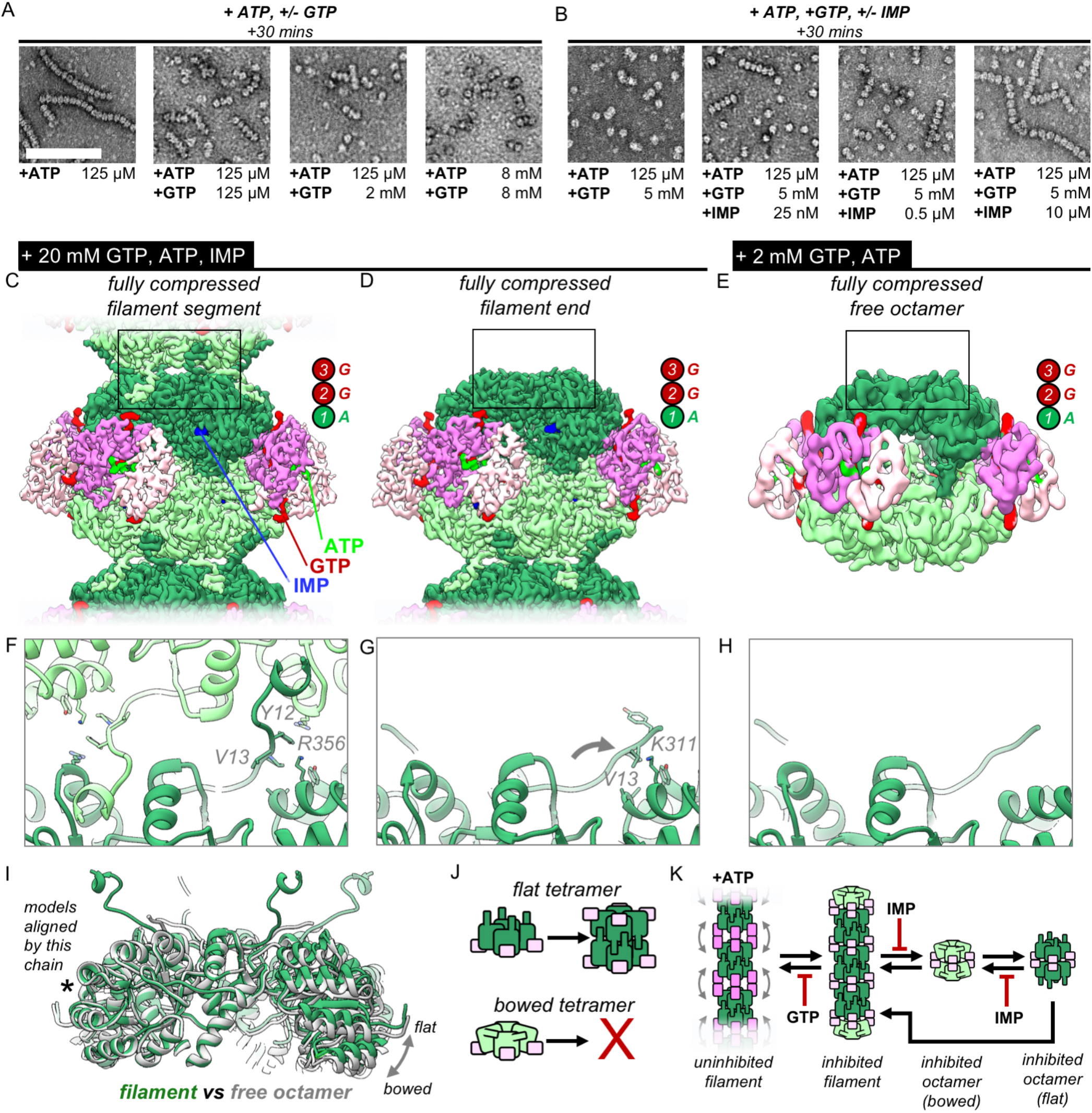
IMP and GTP allosterically modulate filament assembly and disassembly. A) Roughly 2 mM GTP inhibits filament assembly of IMPDH by ATP. Negative stain EM, protein concentration 2 uM. Scale bar 100 nm. B) Roughly 10 uM IMP inhibits filament assembly by GTP. C) Composite cryo-EM density of the GTP/ATP/IMP filament assembly interface and fully compressed filament segment maps. C) Cryo-EM density of the fully compressed filament end map. E) Cryo-EM density of the GTP/ATP non-filament fully compressed free octamer map. F-H) Close-up ribbon views of the assembled and unassembled filament interfaces the maps in A-C. I) Comparison between the tetramer conformations of the “flat” assembled filament interface (green) and the “bowed” unassembled free octamer (gray). J) Cartoon of the relationship between tetramer bowing and filament assembly. K) Model of the regulation of filament assembly by GTP and IMP.

To understand how IMP and GTP allosterically influence filament assembly and disassembly of ATP-bound IMPDH2, we acquired cryo-EM data of the enzyme in two ligand states: 1) ATP/GTP/IMP, and 2) ATP/GTP. To ensure morphological consistency, we sought to saturate the enzyme with GTP. For the ATP/GTP dataset, we used 2 mM GTP, which for both our initial negative stain experiments and cryo-EM preparations resulted in complete disassembly of filaments into free octamers (Fig. S4C). However, under saturating IMP concentrations, 2 mM GTP resulted in filaments that were often bent (Fig. S4D). This should not be possible if GTP were saturating all sites; our structures above suggest bent filaments must contain some extended protomers whose GTP-binding allosteric site 3 is disrupted. For the ATP/GTP/IMP cryo-EM dataset we therefore used a much higher GTP concentration (20 mM), which resulted in filaments with fully compressed segments (Fig. S4E). We explore this apparent difference in GTP affinity in more detail below.

The compressed filament dataset resulted in three high-resolution reconstructions: the consensus filament assembly interface, a fully compressed octameric filament segment, and a fully compressed octameric end segment (Fig. S5A, Table 4). The ATP/GTP/IMP filament assembly interface map (3.0 Å) is identical (backbone RMSD=0.407) to the ATP/IMP/NAD^+^ filament interface (Figs. S5B-C). Classifying the most well-resolved filament segments as before, we obtained a 3.2 Å structure of a fully compressed filament segment (Figs. 4C, S5D-H). The Bateman domains are symmetrically compressed, with ligand density at all three allosteric sites. The active site is well resolved, including the canonical interactions between opposing active site fingers that inhibit catalysis by preventing catalytic dynamics (Buey et al. 2017).

**Table 4.**
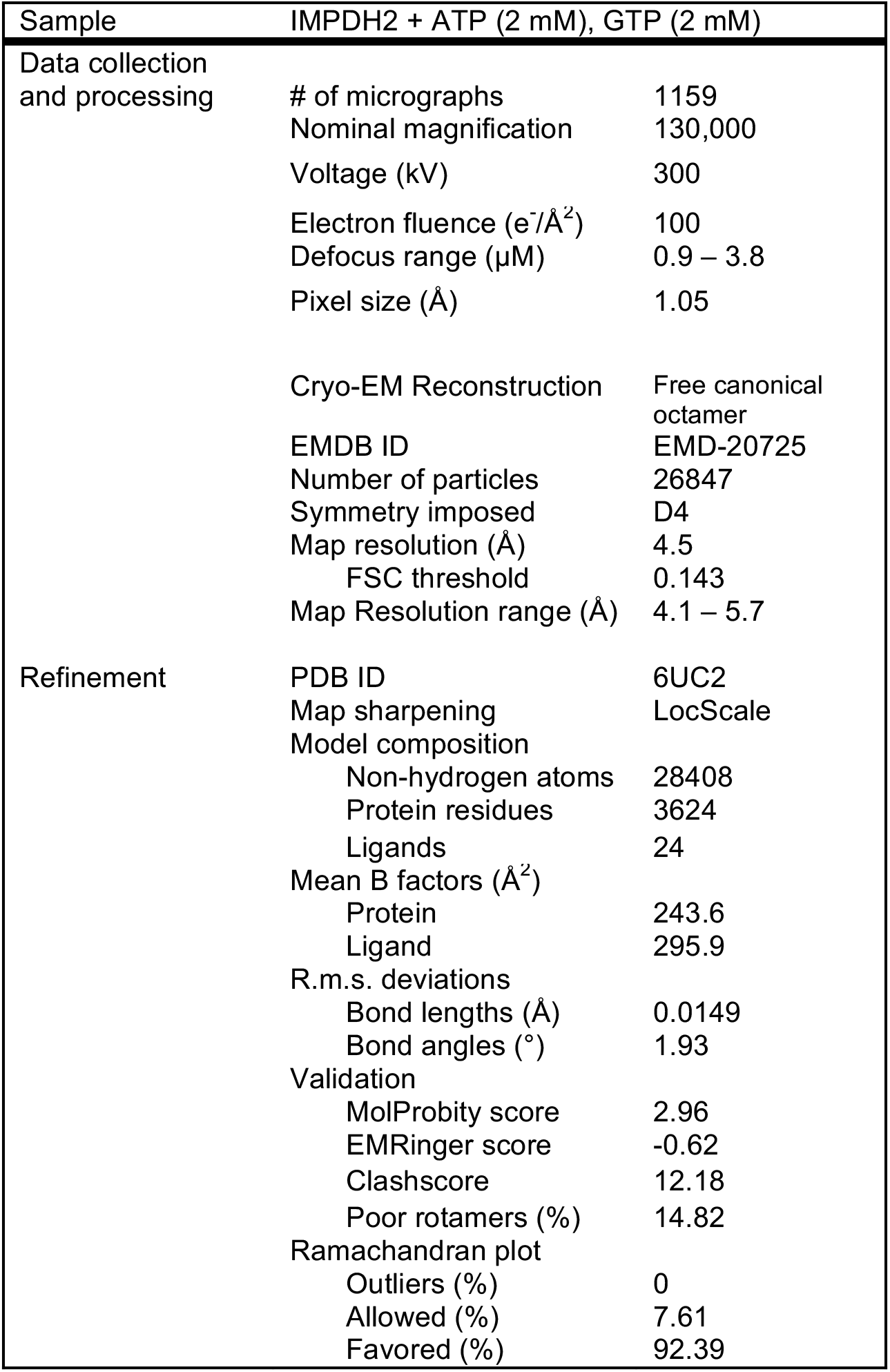
Statistics of cryo-EM data collection, reconstruction and model refinement for the ATP, 2 mM GTP dataset.

Compared to the uninhibited filament datasets, these GTP-saturated, +IMP filaments were shorter in length. As a result, in addition to filament segments, we identified many filament ends: octamers in which one tetramer does not have an assembled interface. The best-resolved structure of these filament ends (3.3 Å) is conformationally similar to the filament segments, being fully compressed, with well-resolved IMP-bound active sites in the inhibited conformation; however, without the filament assembly interface, the N-terminus is only partially resolved (Figs. 4D, S5I-L).

From the free octamer dataset containing ATP/GTP, we used a similar symmetry-expansion and classification strategy to that used for the filament datasets (Fig S6A). This scheme confirmed that virtually all the particles were symmetric, fully compressed free octamers in which both potential filament assembly interfaces are unbound, with some poorly resolved minority classes of larger oligomers (Fig. S6B-C). The best resolved free octamer class reached intermediate resolution (4.5 Å) (Figs. 4E, S6D-F, Table 5). This octamer is conformationally similar (backbone RMSD = 0.978 Å) to a recent higher-resolution crystal structure of IMPDH2 bound to GTP (PDB 6i0o) (Buey et al. 2017; Fernández-Justel et al. 2019). The Bateman domains are fully compressed with three occupied allosteric sites. Without IMP present, the active site loops of the free octamers are disordered, with partial density for the fingers extending across to the opposing tetramer.

**Table 5.**
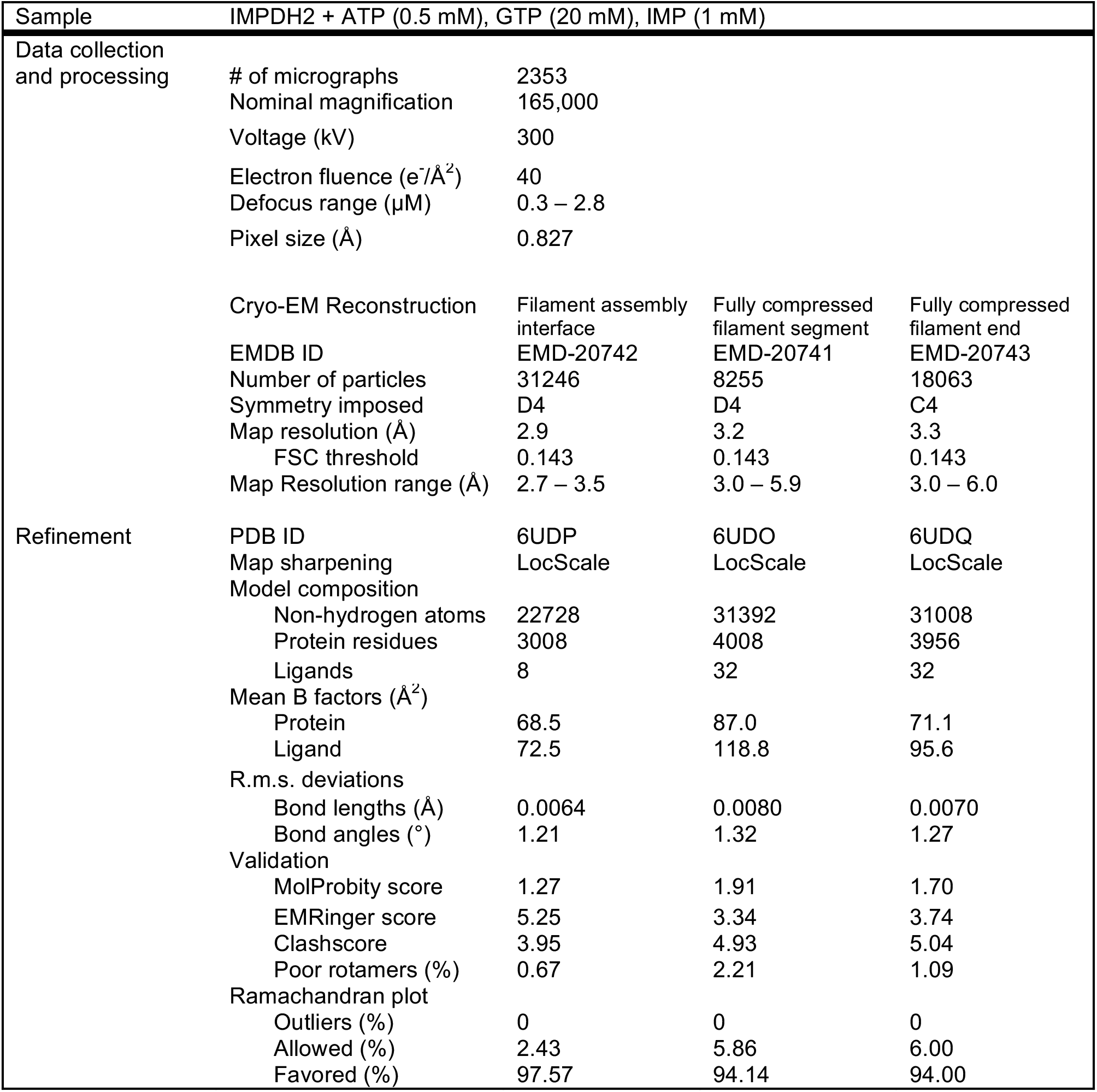
Statistics of cryo-EM data collection, reconstruction and model refinement for the ATP/IMP, 20 mM GTP dataset.

Whether in a filament segment, filament end, or free octamer, the Bateman domains of the three ATP/GTP-bound IMPDH octamer types are fully compressed, with ligand density in all regulatory sites including the critical GTP/GDP-specific third site, which “staples” IMPDH octamers in the fully compressed, inhibited conformation. But we observed two key structural differences that correlated with assembly state: the conformation of the N-terminus, and the relative orientation of protomers within each tetramer. The filament interface of the inhibited segments is unchanged from the uninhibited filaments (Fig. 4F). However, at the free filament ends the N-terminus is only partially unresolved (a.a. 1-11), with the resolvable portion rotated ∼30° degrees compared to the bound interface, such that Val13 inserts into the shallow hydrophobic pocket formed by A307, A308, and Lys311 of the neighboring protomer, very similar to its position in the GTP-bound free octamer crystal structure (PDB 6i0o) (Fig. 4G) (Fernández-Justel et al. 2019). The resolution of the free octamer structure precludes side chain placement, but the backbone density of the unbound N-terminus is in the same position, indicating that free filament ends and free octamers are in the same conformation (Fig. 4H).

We noticed a striking difference between the conformation of the catalytic core assembly in free versus assembled states. Each protomer of the free octamer and at filament ends is tilted ∼5° relative to the four-fold symmetry axis, such that the tetramer becomes more “bowed” than the “flat” tetramers at filament assembly interfaces (Fig. 4I, Video S1). The tetramers found in the GTP-bound free octamer crystal structure (PDB 6i0o) are ina similar bowed conformation (Fernández-Justel et al. 2019). The bowed tetramer conformation and the filament assembly interface are mutually exclusive. When the N-terminus conformation from the free octamer is modeled into the flat tetramer of the filament assembly interface, V13 is not correctly positioned to bind the A307/A308/K311 of the neighboring protomer, and instead clashes with K311 and A307 (Fig. S7A). Thus the flat conformation promotes release of the N-termini from their binding sites on neighboring protomers, freeing them to rotate into the conformation seen in the assembled filament interface. The reciprocal operation is also not possible; for a bowed tetramer modelled with the N-terminus conformation from the filaments, only one protomer at a time can form the assembled interface (Fig. S7B). The other N-termini are out of position, with significant separation of the critical residues Y12 and R356 as well as multiple steric clashes. Thus bowed tetramers cannot form the IMPDH2 filament assembly interface, and tetramer flattening is a necessary precondition for IMPDH2 filament assembly (Fig. 4J).

These three structures of inhibited IMPDH2 conformations provide a model by which filament assembly is influenced by IMP and GTP through tetramer bowing and flattening (Fig 4K). In the absence of substrate, GTP induces both compression of filament segments and tetramer bowing, with the latter resulting in filament disassembly. But when IMP is bound, the disordered active site loops become ordered and rigid, buttressing intra-tetramer contacts as well as forming a pseudo-beta-barrel between opposing tetramers in the GTP-bound state. These increased contacts work to resist tetramer bowing and more readily sample the flat tetramer conformation, which promotes, and is stabilized by, the filament assembly interface. When IMP levels are low, GTP promotes filament disassembly, but high IMP levels shift the equilibrium towards filament assembly.

### IMPDH2 filaments resist GTP inhibition

Based on this model, during quiescence, when the salvage pathways supply ample GTP and IMP production is downregulated, intracellular IMPDH2 will be in the fully compressed, fully inhibited, ATP/GTP-bound free octamer state. Without increased IMP production, guanine depletion will result in transient octamer extension and filament assembly, both of which will reverse as the resulting increase in IMPDH flux restores guanine levels by diverting substrate from the parallel de novo adenine pathway, mirroring the extension/compression behavior of non-filament assembling homologues (Fernández-Justel et al. 2019; Buey et al. 2017). Recently, it was shown that in vivo IMPDH assembly is promoted by increases in intracellular IMP (Keppeke et al. 2018). We therefore reasoned that the primary regulatory function of filament assembly may apply only in the proliferative state, when intracellular IMP levels are upregulated, preventing filament disassembly by GTP.

To probe whether filament assembly influences the regulatory effects of GTP on IMPDH2 activity we compared enzyme kinetics of the wildtype enzyme with the non-assembling mutant Y12A. We measured the GTP dose-response of IMPDH2 pre-incubated with varying levels of ATP, and found that wild-type enzyme is less sensitive to GTP inhibition, compared with Y12A (Figs. S8A-C). Depending on ATP concentration, the apparent GTP IC50 of the wild-type was roughly two-fold lower than for Y12A. At higher ATP levels, the apparent GTP IC50 of WT and the non-assembly mutant both increase. We attribute this to competition between ATP and GTP at the first and second Bateman sites, suggesting that independent of filament assembly, GTP inhibition of IMPDH2 is affected by ATP. Notably, the range of GTP in which the substrate-saturated filaments resist GTP inhibition is within the upper range of in vivo concentrations (Traut 1994). One function, then, of IMPDH2 filaments is to resist inhibition by GTP.

To correlate structural differences in IMPDH2 filaments as a function of GTP concentration, we collected negative stain EM data directly from three reaction volumes used for the GTP inhibition experiments, corresponding to uninhibited filaments (no GTP), ∼10% inhibited filaments (2.5 mM GTP), and fully inhibited filaments (20 mM GTP) (Fig. 5A). As in the cryo-EM datasets, we found that uninhibited filaments were often extended, and fully inhibited filaments were universally compressed. However, we noted that the partially inhibited filaments contained a more heterogeneous mix of extended, bent, and compressed segments.

**Figure 5.**
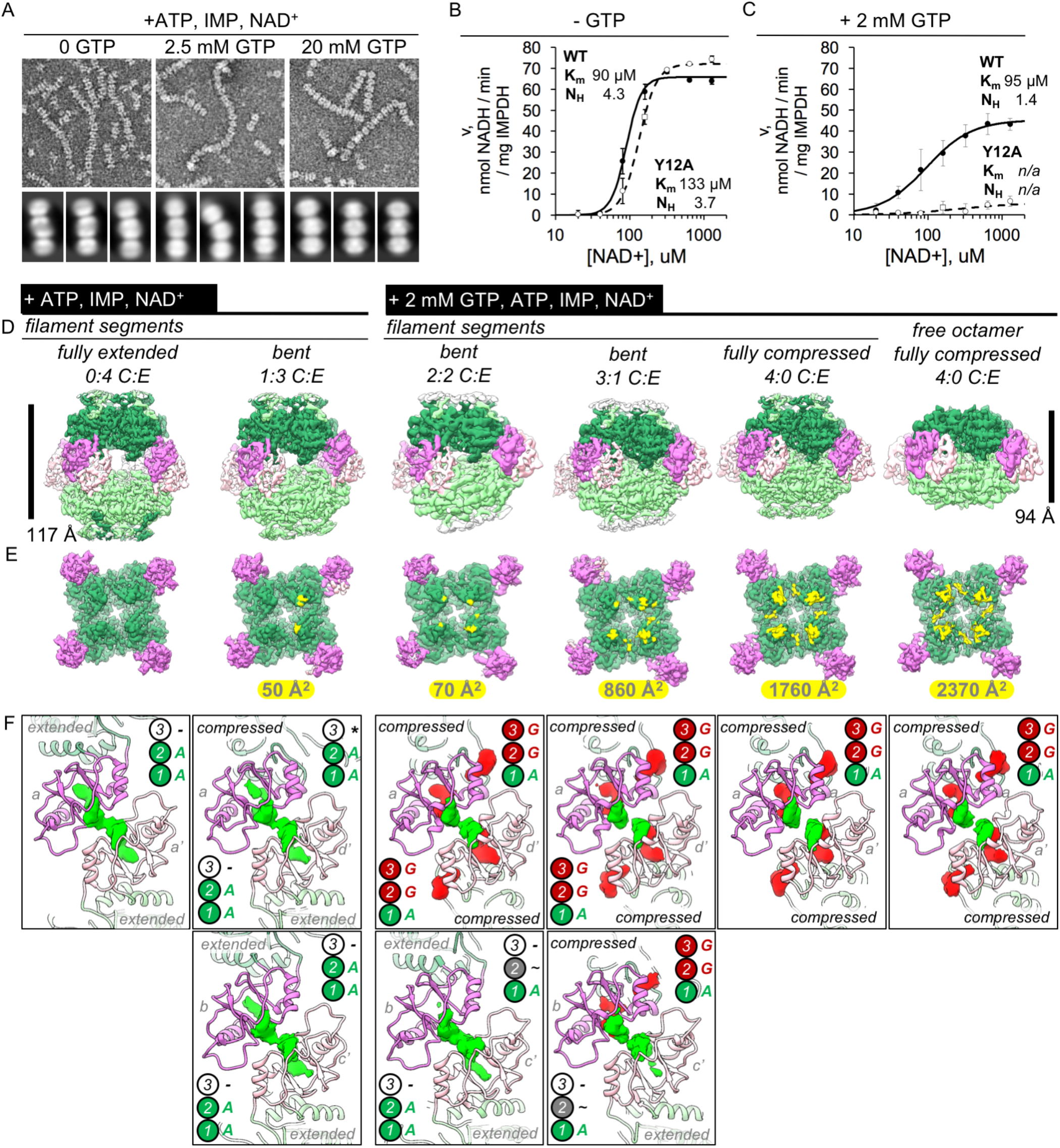
IMPDH2 filaments resist GTP inhibition by promoting bent octamer conformations that separate opposing active sites. A) Negatively stained EM of uninhibited (left), partially inhibited (center), and fully inhibited (right) IMPDH2. Representative micrographs and reference free 2D class averages. B) NAD^+^ saturation curves of uninhibited WT IMPDH2 (solid line), and the non-assembly mutant Y12A (dashed line). Reactions performed with 0.5 mM ATP, 1 mM IMP. C) NAD^+^ saturation curves of WT filaments treated with 2 mM GTP. D) Six cryo-EM maps from two datasets (uninhibited ATP/IMP/NAD^+^ and partially inhibited ATP/IMP/NAD^+^/[2mM]GTP) exhibiting a range of Bateman domain conformations. E) A view of a single tetramer from the inside of each octamer. The lighter colored tetramer from panel A is hidden, with the surface area buried between tetramer active sites colored in yellow, with the indicated total buried surface area. F) Corresponding views of Bateman domain conformations. Protein displayed as ribbon, with the two interacting Bateman domains colored orchid and light pink. Cryo-EM density for non-protein ligand densities is colored green and red, for ATP and GTP, respectively. Symmetry identities labelled with gray letters. In the extended conformation, allosteric site 3 is distorted, and does not bind ligands (black dashes). Allosteric site 3 is formed in compressed protomers, but in the absence of guanine nucleotides it remains unoccupied (black asterisk). In the extended protomers of some bent octamers, is is possbile that allosteric site 2 is only partially occupied (black tilde).

Next, we compared NAD^+^ kinetics of uninhibited (0 mM GTP) and partially inhibited (2 mM GTP) filaments with saturating IMP concentrations (Figs. 5B-C). As expected, in the absence of GTP WT and the non-assembling mutant have similar apparent Michelis-Menton kinetics (Anthony et al. 2017). Inhibition of WT by this concentration of GTP is partial and non-competitive. In contrast, the non-assembling mutant is strongly inhibited. In the absence of saturating IMP, these same GTP levels result in complete octamer compression and filament disassembly, providing a possible mechanism by which filament assembly alters GTP inhibition: by resisting the fully compressed state.

To better understand this partially inhibited state, we collected cryo-EM data of IMPDH2 with saturating substrates, 0.5 mM ATP, and 2 mM GTP (Figs. S8D). Under these conditions we observed the non-assembly mutant Y12A was 84% inhibited but WT enzyme was only 16% inhibited. We resolved to high resolution structures of not only the filament assembly interface and several distinct filament segments, but surprisingly, two different free octamers; a canonical free octamer (a dimer of tetramers bound by Bateman domains), and also a small class of free interfacial octamers (a dimer of tetramers bound by the filament assembly interface) (Figs. S8E, S9A, Table 6).

**Table 6.**
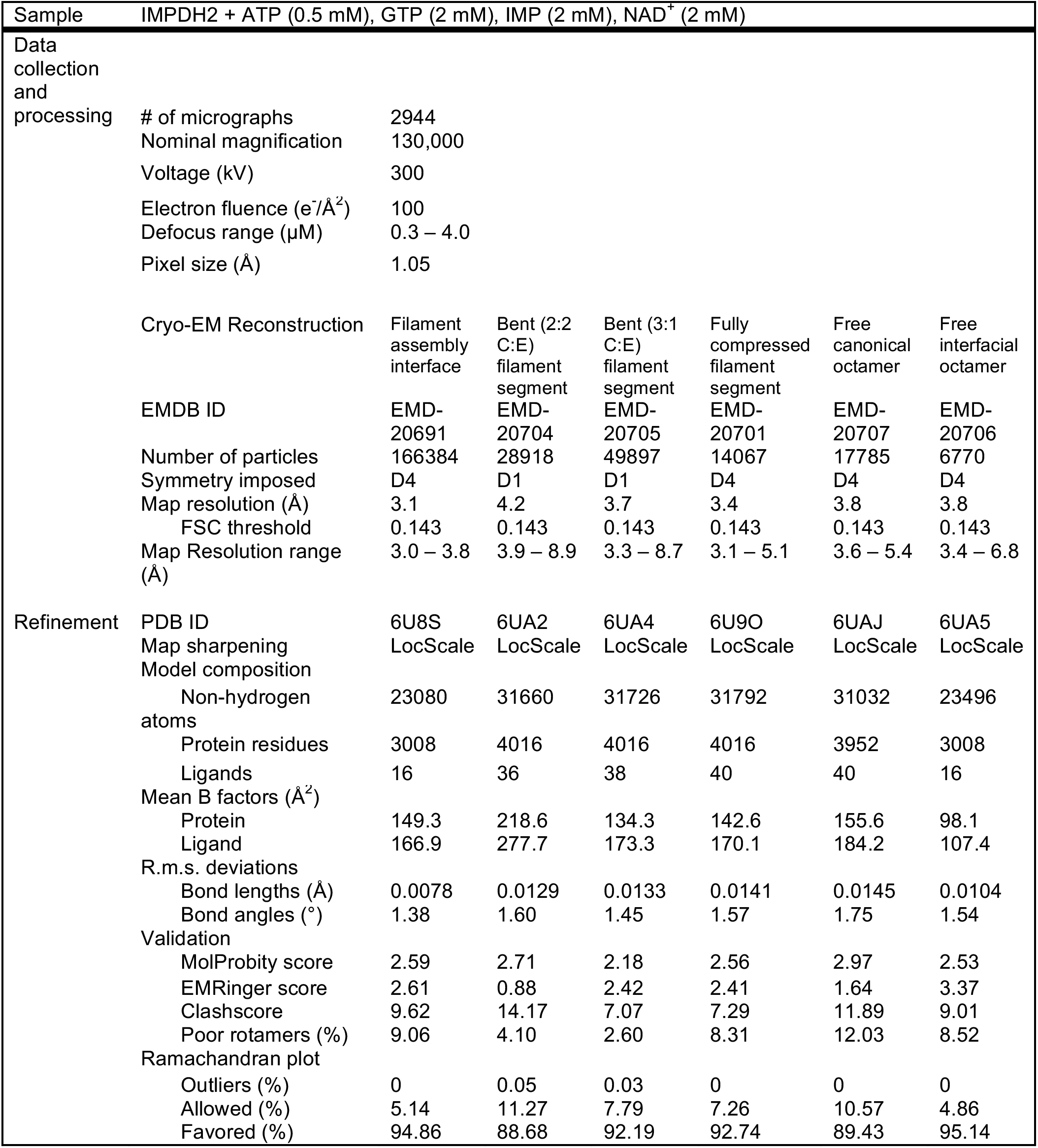
Statistics of cryo-EM data collection, reconstruction and model refinement for the ATP/IMP/NAD^+^, 2 mM GTP dataset.

The 3.1 Å resolution consensus assembly interface map is identical to the uninhibited and fully inhibited consensus interfaces, including a well-resolved active site with strong density for both substrates; as with the ATP/IMP/NAD+ structures these filaments are actively turning over and we have likely captured many states, which we have modelled simply as IMP/NAD^+^ (Figs S9B-C, S12A). As expected, the filament segments exhibited a range of conformations (Fig. S9D). The best resolved of these were a fully compressed filament segment at 3.4 Å resolution, and two bent conformations at 4.2 and 3.7 Å resolution (Figs. S9E-M). The fully compressed filament segment is identical to the corresponding structure from the inhibited (20 mM GTP) filament dataset, with all Bateman binding sites occupied (Fig. S12B). As with the bent conformation from the ATP/IMP/NAD^+^ dataset, the two bent filament segments are each two-fold symmetric octamers of asymmetric tetramers containing different proportions of extended or compressed protomers (Figs. S12C-D). The asymmetric unit from one of these bent segments is a tetramer with two compressed and two extended protomers, and the other has three compressed and one extended. For each of the compressed protomers, there is clear ligand density bound at Bateman Site 3; for the extended protomers this site is unformed and empty.

Unlike the mixture of extended and compressed protomers we observed in the filaments, the free canonical octamers were universally compressed (Fig. S10A-C). As with the ATP/GTP free octamer, the best resolved canonical free octamer class (3.8 Å) is fully compressed and bowed, despite having both substrate sites occupied (Figs. S10D-F, S12E). Thus, even in the IMP-bound state, the compressed/flat conformation is less preferred, unless stabilized by the filament assembly interface. The small class of free interfacial octamers (3.8 Å) was no different from the filament interface maps, except that the Bateman domains are completely unresolved, indicating that without the stabilization provided by Bateman domain dimerization, these regions are highly flexible (Figs. S11A-F, S12F). The observation of dramatically different structural ensembles in filament-bound and free IMPDH2, in the presence of identical ligand concentrations, explains the role of filament assembly in resisting compression and GTP inhibition.

### Filament-specific IMPDH2 conformations reduce GTP affinity and promote activity

From our different cryo-EM datasets combined, we have now determined structures of canonical IMPDH2 octamers bound to allosteric effectors and both substrates, in six distinct conformations (Fig. 5D). From the uninhibited ATP/IMP/NAD^+^ dataset we resolved a fully extended filament segment, and a bent segment in which for each tetramer, 3 protomers were extended and 1 was compressed. From the partially inhibited GTP/ATP/IMP/NAD^+^ dataset, we resolved 2:2, 3:1, and fully compressed filament segments, and a fully compressed free octamer. These five structures provide a mechanism by which the Bateman domain extension of discrete protomers promoted by filament assembly resists GTP inhibition (Fig 5E). Depending on the degree of extension/bending/compression, filament segments exhibit a range of increasingly stronger interactions between opposing catalytic domains. For the fully extended segments, there are no interactions, and the flexible active loops are able to perform the complex conformational changes necessary for catalysis (Hedstrom 2009; Buey et al. 2017). Going through the progressively more compressed bent states, there is progressively greater surface area buried by a series of distinct contacts between the opposing active site loops, until the fully compressed filament segment. For the active sites that make these contacts, activity is likely impaired because the active site loops are constrained. However, the presence of some unconstrained active sites in the bent filament segments means that these protomers are catalytically active. This effect is not cooperative within the octamers, and even a single extended protomer is sufficient to reduce inhibitory active site contacts.

The Bateman domain ligand occupancy of these five filament segment conformations varies significantly. Given the resolution range of these structures (3-4 Å) it is not possible to unequivocally distinguish ATP from GTP, and we have assigned ligand identity according to previous structures of a bacterial IMPDH bound to ATP (PDB 4dqw) (Labesse et al. 2013) and a fungal IMPDH bound to ADP and GTP (PDB 5tc3) (Buey et al. 2017) (Fig. S14C). These were chosen due to conformational similarity of the Bateman domains of these structures to our ATP/IMP/NAD^+^ fully extended and ATP/IMP/GTP fully compressed human filament cryo-EM structures; backbone RMSDs were 1.163 and 0.856 Å, respectively (for residues corresponding to human IMPDH2 residues 110-244). As previously described, in both the fully extended and 1:3 compressed:extended segments there is clear ligand density at Bateman sites 1 and 2 (Fig. 5F). For the single compressed protomer in the latter, site 3 is formed but due to the absence of GTP in that dataset it is unoccupied. In the partially inhibited filament dataset, for which the buffer contained both ATP and GTP, we see a greater variation in Bateman ligand occupancy. For the 2:2 compressed:extended segment, there is full occupancy at sites 1, 2, and 3 in the compressed protomers, however for the two extended protomers there is strong density at site 1 and partial density at site 2. The 3:1 compressed:extended segment is qualitatively similar; the three compressed protomers possess full ligand occupancy but for the one extended protomer site 2 has only partial density. For both the fully compressed filament segment and the free octamer there is full ligand density. Thus filament assembly promotes the extended state of individual protomers, which reduces overall GTP affinity due to disruption of site 3.

## Discussion

Understanding the complex ways cells regulate IMPDH activity to efficiently maintain spatiotemporal control of nucleotide levels in response to varying demand has direct implications for human health. IMPDH activity is upregulated to increase guanine levels in proliferating tissues like tumors and regenerating liver (Tressler et al. 1994; Yalowitz & Jayaram 2000; Huang et al. 2018; He et al. 2018; Nagai et al. 1991). IMPDH plays a particularly important role in the immune response, where T-cell activation is dependant on increased production of purine nucleotides, and is associated with IMPDH filament assembly (Gu et al. 2000; Zimmermann et al. 1998; Duong-Ly et al. 2018; Calise et al. 2018). As a result, IMPDH is the target of several drugs used in immunosuppressive treatment of both autoimmune disease and organ transplant rejection, and is considered a promising target for antineoplastic agents (Shu & Nair 2008; Liao et al. 2017; Bergan et al. 2016).

Assembly and disassembly of IMPDH into filaments has been observed in healthy proliferative cells, and in cancer cells (Keppeke et al. 2018; Wolfe et al. 2019). IMPDH filaments reversibly assemble in stimulated T-cells as they transition to a proliferative state, in a mechanism dependent on multiple metabolic signaling pathways and on the levels of guanine nucleotides (Duong-Ly et al. 2018; Calise et al. 2018). Despite the importance of understanding human IMPDH regulation, until now the regulatory role of IMPDH filament assembly has not been explored at the structural level.

We propose a model that describes the regulatory role of human IMPDH2 filament assembly, in which assembly reduces feedback inhibition of enzyme activity in a substrate-dependant manner, increasing flux through the de novo guanine nucleotide synthesis pathway in response to proliferative signalling. In the absence of either IMP or guanine ligands, IMPDH2 is conformationally dynamic (Fig. 6A). Apo IMPDH2 forms stable tetramers, which freely sample both the “bowed” and “flat” tetramer conformations, with the latter resulting in release of the N-terminus and assembly into stable interfacial octamers. Adenine nucleotides bind with high affinity to the Bateman domain, resulting in stable filaments in which the active site loops remain unconstrained and the enzyme is active. The ATP concentration required to induce filament assembly *in vitro* is far below the expected *in vivo* levels; we thus predict the apo state to be rare in human cells (Traut 1994). Without bound guanine nucleotides, the Bateman domains of individual protomers freely compress and extend (Fig. 6B). However, the flat tetramer conformation found in assembled IMPDH2 filaments is resistant to full compression of octameric filament segments. Binding of GTP to the Bateman domain stabilizes the compressed state, leading to lattice strain that is relieved by disassembly of the filament interface and tetramer bowing. In this way high intracellular guanine levels disassemble IMPDH2 filaments into stable free octamers whose activity is inhibited. Binding of IMP to the active site stabilizes the flexible active site loops, and saturation with both IMP and GTP results in compressed filaments in which the active sites of opposing tetramers interlock into a stable network. This rigidifies the octameric filament segment, which now resists the lattice strain brought on by compression, blocking GTP-induced filament disassembly.

**Figure 6.**
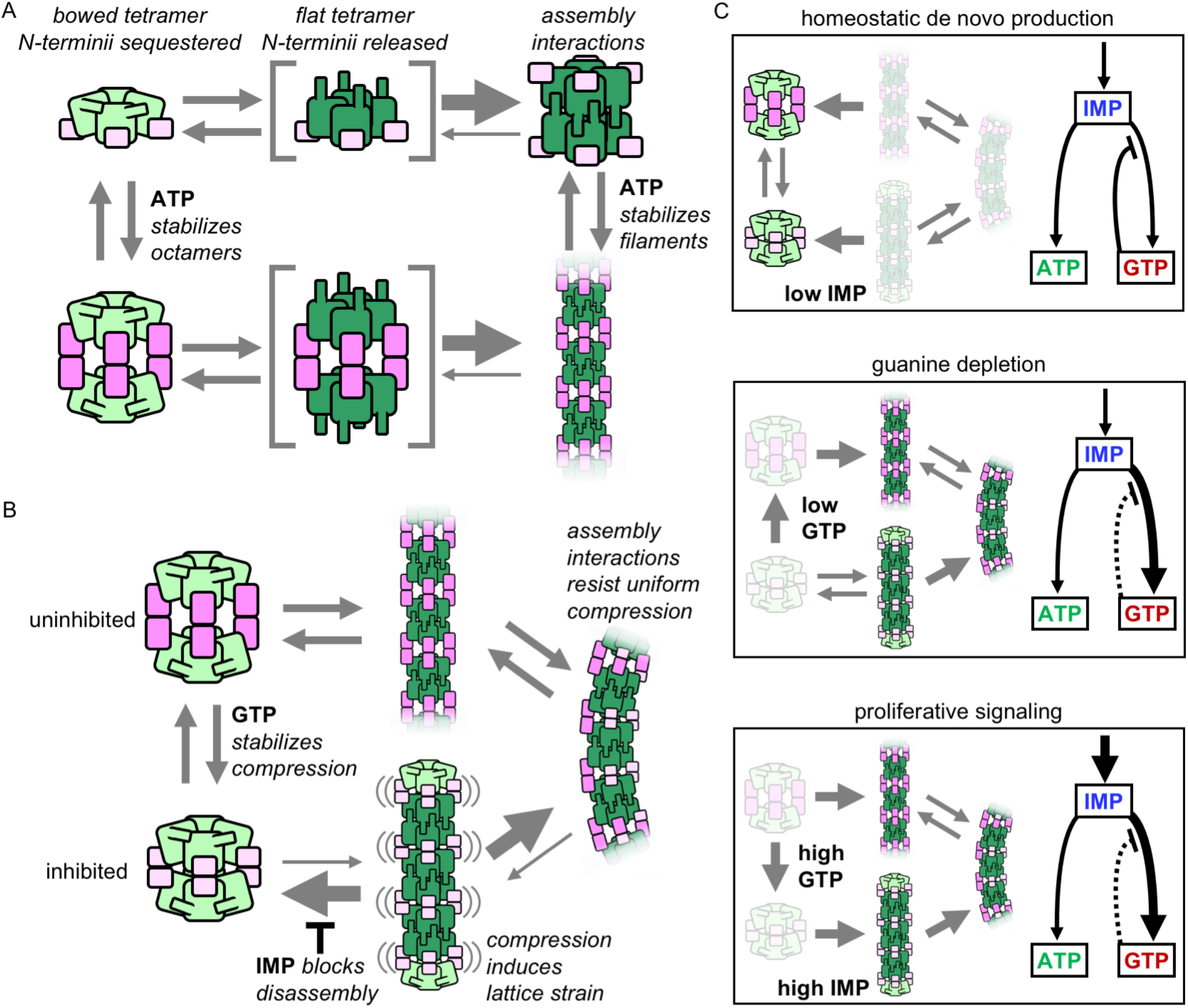
Model of IMPDH2 assembly and filaments’ role in guanine nucleotide regulation. A) Filament assemble when octamer interactions are stabilized by ATP binding to the regulatory domain (pink), and the N-terminal residues (blue) are released by flattening of the catalytic tetramer (green). Filament assembly interactions stabilize the flat conformation. B) GTP binding stabilizes an inhibited, compressed conformation. Filament is less sensitive to GTP-induced compression, maintaining a population of octamers in mixed activity states. Filaments in the fully compressed GTP-bound state are strained, which promotes disassembly that is inhibited by substrate IMP binding. C) The equilibrium in (B) explains the different cellular conditions in which IMPDH polymerization occurs. i) Under homeostatic conditions IMPDH2 is dispersed and activity is regulated by GTP binding to octamers, which balances low levels of de novo synthesis between adenine and guanine pathways. ii) When guanine nucleotides are depleted the equilibrium shifts toward filaments. iii) Proliferative signaling can directly shift the equilibrium toward filaments, where higher flux is maintained through the guanine pathway under elevated GTP concentrations due to reduced sensitivity of the filaments to GTP inhibition (dashed line).

By balancing these complex conformational dynamics, cells fine-tune feedback inhibition of the enzyme, consistent with the states in which IMPDH filaments are observed in cells (Fig. 6C). Under homeostatic conditions, the salvage pathways provide sufficient purine nucleotides, and IMP production is low. Under these conditions IMPDH2 is bound to both adenine and guanine nucleotides, but not IMP, forming free octamers rather than filaments. In vivo, filaments are typically not observed in quiescent cells; rather, as in our model, their assembly is associated with increased intracellular IMP and decreased intracellular GTP (Schiavon et al. 2018; Keppeke et al. 2018; Calise et al. 2014; Juda et al. 2014). Depletion of guanine allows Bateman domain extension, and transient assembly into enzymatically active filaments. However, upon restoration of guanine levels, these active filaments disassemble into compressed free octamers, mirroring the known feedback inhibition behavior of non-filament forming IMPDH homologues (Buey, Ledesma-Amaro, Velázquez-Campoy, et al. 2015; Buey et al. 2017; Fernández-Justel et al. 2019).

The key difference is the substrate dependance of IMPDH2 filament assembly. IMP-saturated IMPDH2 filaments resist both disassembly and the fully compressed, inhibited state. Even at elevated guanine nucleotide levels, these filaments retain a proportion of uninhibited active sites. This allows the cell to modulate enzyme activity to balance levels of product and substrate in response to metabolic demand, which can vary significantly depending on cell type and cell cycle stage. Our data strongly support the idea that IMPDH2 filament assembly serves to elevate intracellular guanine nucleotide levels during the proliferative state by resisting feedback inhibition. Production of IMP is upregulated in response to overall purine demand, as well as in response to proliferative signalling via the mechanistic target of rapamycin (mTOR) (Smith 1998; Ben-Sahra et al. 2016). Inhibition of mTOR reverses IMPDH filament assembly in activated mouse T-cells, as well as proliferating mouse liver cells (Duong-Ly et al. 2018; Chang et al. 2015). Thus the dependence of filament assembly on IMP levels provides an avenue for regulation of assembly through established proliferative signalling pathways.

Many metabolic enzymes form filamentous polymers in cells in response to changes in metabolic state (Narayanaswamy et al. 2009; Ingerson-Mahar et al. 2010; Noree et al. 2010; O’Connell et al. 2012; Zhao et al. 2013; Petrovska et al. 2014; Shen et al. 2016; Saad et al. 2017). Most of these metabolic filaments are assembled from important regulatory enzymes, which suggests polymerization may play a role in modulating flux through these pathways. Recently, in just a few cases, structural and biochemical studies have provided insight into the functional consequences of enzyme filament assembly, suggesting that one role of polymers is to regulate activity by locking enzymes into active or inactive conformations through assembly contacts (Hunkeler et al. 2018; Barry et al. 2014; Lynch et al. 2017; Webb et al. 2018; Stoddard et al. 2019). Thus, it was surprising when our initial characterization of IMPDH2 filaments showed that assembly did not affect enzymatic activity or the ability to switch between active or inactive conformations (Anthony et al. 2017). Instead, as we have shown here, IMPDH2 filaments fine-tune the the allosteric response by reducing affinity for inhibitory downstream products. This represents a new way in which enzyme assemblies can modulate flux through metabolic pathways, providing an additional layer of regulatory control on top of existing transcriptional, post-translational, and allosteric regulation.

## Author Contributions

Conceptualization, M.C.J. and J.M.K.; Methodology, M.C.J. and J.M.K.; Validation, M.C.J; Formal Analysis, M.C.J.; Investigation, M.C.J.; Resources, J.M.K.; Data Curation, M.C.J.; Writing – Original Draft, M.C.J. and J.M.K.; Writing – Review & Editing, M.C.J. and J.M.K.; Visualization, M.C.J.; Supervision, J.M.K.; Project Administration, J.M.K.; Funding Acquisition, J.M.K.

## Acknowledgements

The authors thank the Arnold and Mabel Beckman Cryo-EM Center at the University of Washington for electron microscope use. We are grateful to J. Quispe for technical support and advice. We thank A. Burrell, G. Cai, A. Horowitz, K. Hvorecny, and E. Lynch for valuable feedback. This work was supported by the US National Institutes of Health (R01 GM118396 to J.M.K.).

## Declaration of Interests

The authors declare no competing interests.

## Supplemental Figures

**Figure S1.**
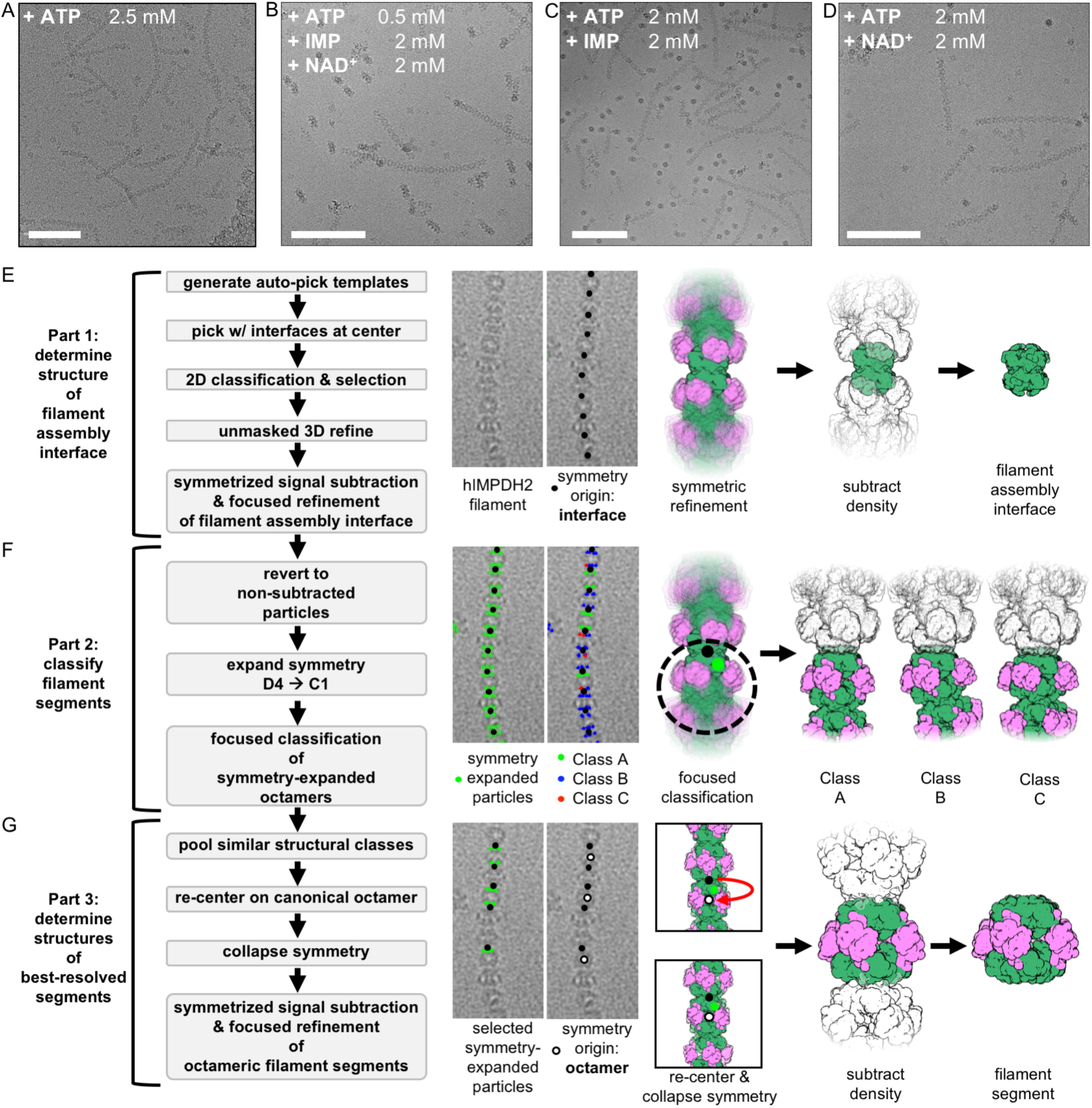
A cryo-EM image processing workflow to for structure determination of flexible IMPDH2 filaments. Related to Figure 2. A-D) Representative cryo-EM micrographs of IMPDH2 treated with ATP (A), ATP and both substrates (B), ATP and IMP (C), or ATP and NAD^+^ (D). Full datasets contained 480, 2169, 2289, and 2178 micrographs, respectively. Scale bars 100 nm. E) Template-based picking, unmasked refinement, density subtraction, and masked refinement results in a reconstruction of the eight symmetrically arranged catalytic domains that make up the filament assembly interface. F) Reverting to the un-subtracted particles, expanding the D4 symmetry, and classifying without alignment using a mask including a single filament segment identifies different segment conformations. G) The best resolved map of each filament segment class was obtained by pooling similar classes, re-extracting and re-centering the refinement from the assembly interface onto to the canonical octamer, collapsing the symmetry expansion by deleting all Euler angle priors and removing overlapping particles, and re-refining from scratch, with additional classification and application of point-group symmetry resulting in further improvements in resolution.

**Figure S2.**
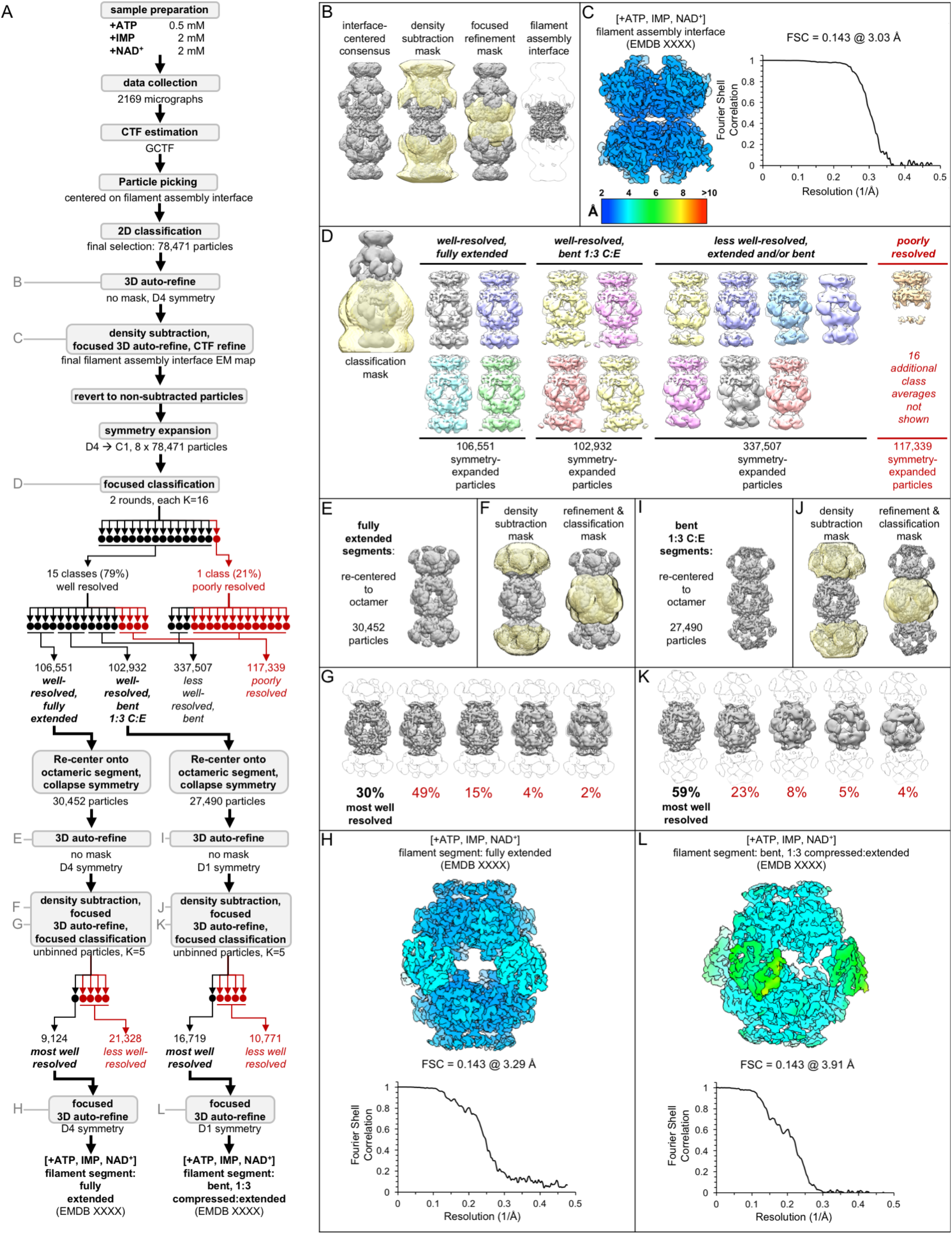
Image processing of the IMPDH2 +ATP, IMP, NAD^+^ cryo-EM dataset. Related to Table 1. A) Flow chart summarizing data processing strategy. B) Density subtraction and focused refinement of the consensus filament assembly interface. C) Local resolution estimation and FSC curve (via relion postprocessing) for the ATP/IMP/NAD+ consensus filament assembly interface. D) Final class averages from symmetry expanded classification of filament segments. E) Unmasked refinement from all fully extended segments, pooled and re-centered. F) Masks used for continued processing of fully extended segments. G) Final classification of the best-resolved fully extended filament segment class H) Local resolution estimation and FSC curve for the ATP/IMP/NAD+ fully extended filament segment I-L) Same as E-H, but for the best-resolved ATP/IMP/NAD+ bent filament segment

**Figure S3.**
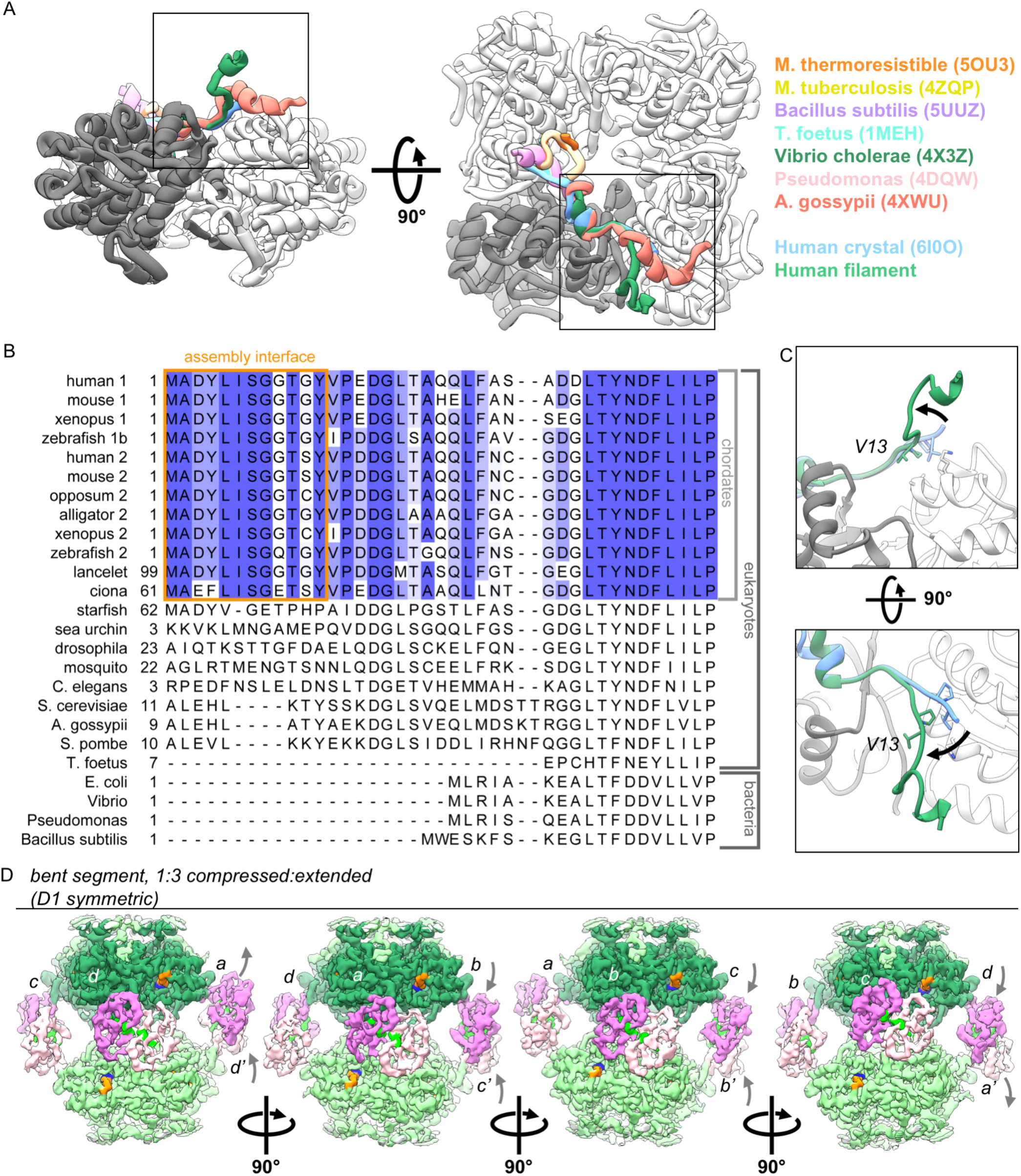
The vertebrate-specific N-terminus mediates IMPDH2 assembly of ATP-bound IMPDH2 filaments, in which individual protomers can extend or compress freely. Related to Figure 3. A)The conformation of the N-terminus seen in assembled filaments of human IMPDH2 is unique among solved IMPDH structures, including other human structures. Boxed regions correspond to views in panel C. B) Sequence alignment of human IMPDH1 (human 1) and IMPDH2 (human 2) and other IMPDH homologues. C) Comparison of N-terminus conformations from cryo-EM of assembled filament (green) and published crystallized IMPDH2 (blue, PDB ID 6I0O). F) Rotated views of the cryo-EM density for the best resolved ATP/IMP/NAD^+^ bent structure, colored as in Fig 2. The asymmetric unit is a tetramer, and each of the four chains can be viewed by rotating incrementally by 90 degrees. Gray letters and arrows indicate chain symmetry mates and Bateman domain conformations.

**Figure S4.**
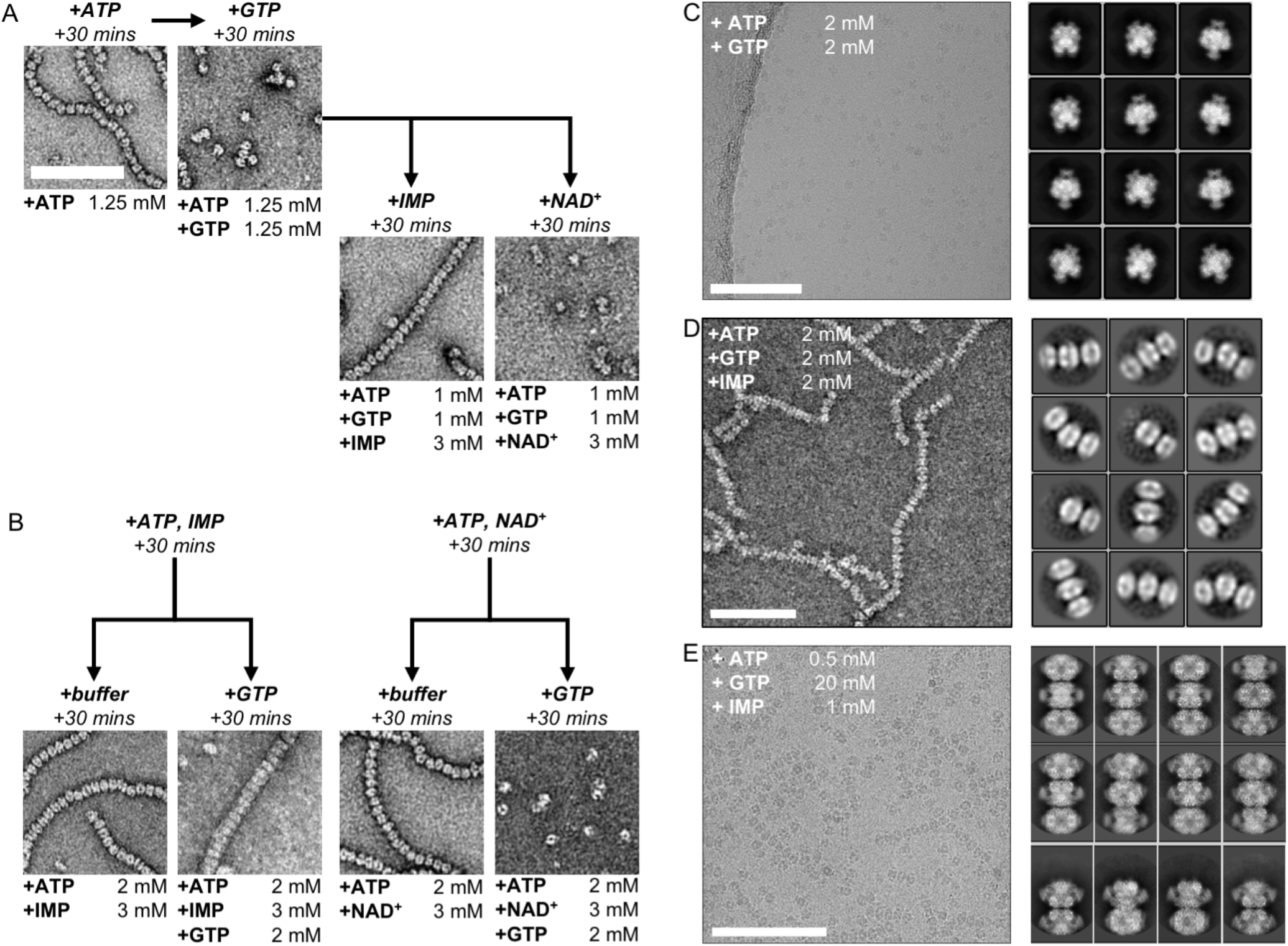
Electron microscopy of human IMPDH2 treated with ATP, GTP, IMP, and NAD^+^. Related to Figure 4. A) IMP, but not NAD+, promotes re-assembly of GTP-disassembled filaments. Reagents added sequentially, with 30 minut room-temperature incubation steps between. B) IMP, but not NAD+, protects against disassembly of filaments by GTP. Reagents added sequentially, with 30 minut room-temperature incubation steps between. C) Cryo-EM of IMPDH2 treated with ATP and 2 mM GTP. Representative micrograph (of 1159) and 2D class averages. D) Negative stain EM f IMPDH2 treated with ATP, IMP, and 2 mM GTP. Representative micrograph and 2D class averages. E) Cryo-EM of IMPDH2 treated with ATP and 2 mM GTP. Representative micrograph (of 2248), and 2D class averages. All scale bars 100 nm.

**Figure S5.**
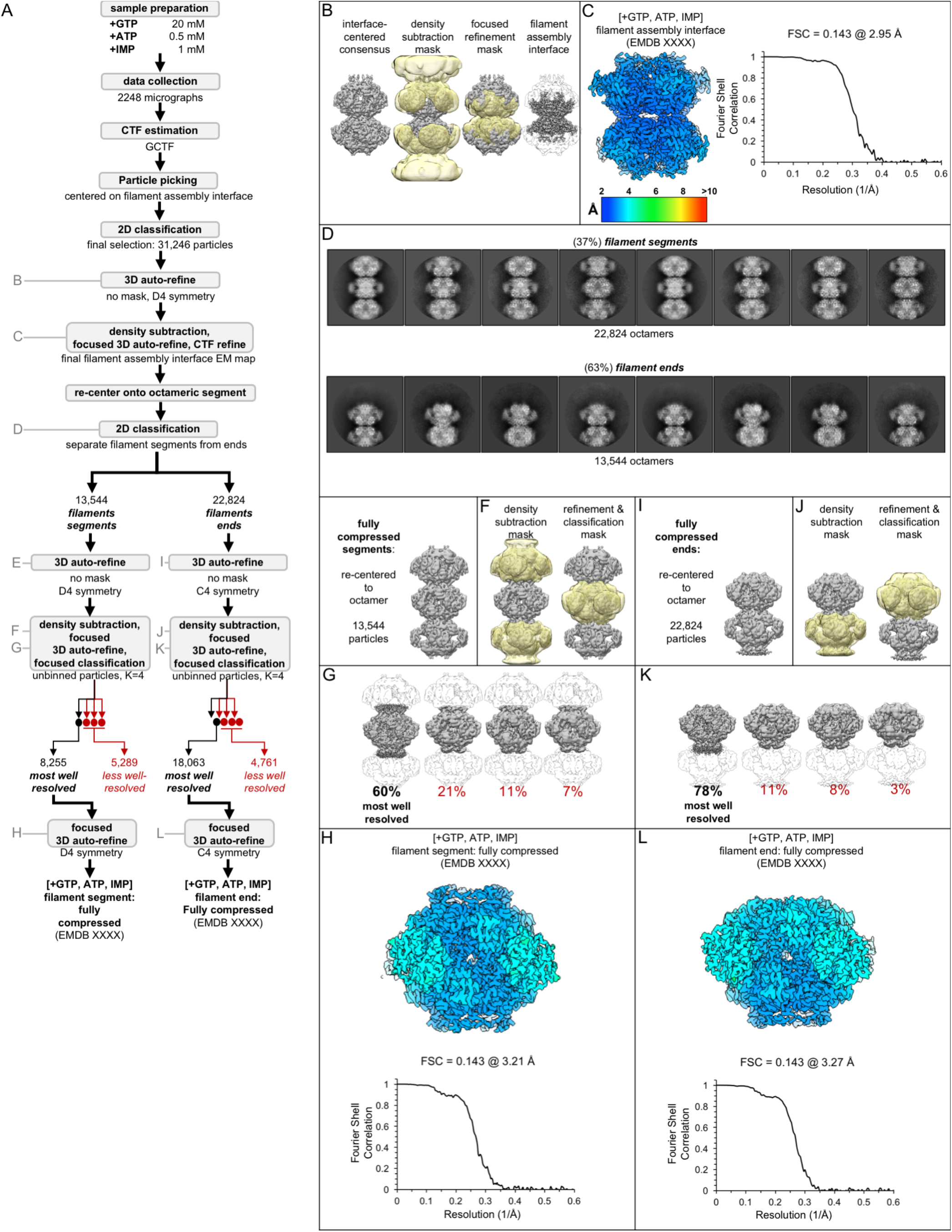
Image processing of the IMPDH2 +ATP, IMP, 20 mM GTP cryo-EM dataset. Related to Table 5. A) Flow chart summarizing data processing strategy. B) Density subtraction and focused refinement of the consensus filament assembly interface. C) Local resolution estimation and FSC curve (via relion postprocessing) for the ATP/IMP/NAD+ consensus filament assembly interface. D) 2D classification to separate filament segments and filament ends. Representative 2D class averages. E) Unmasked refinement from all fully compressed segments, pooled and recentered. F) Masks used for continued processing of fully compressed segments. G) Final classification of the best-resolved fully compressed filament segment class H) Local resolution estimation and FSC curve for the ATP/IMP/[20 mM]GTP fully compressed filament segment I-L) Same as E-H, but for the best-resolved ATP/IMP/[20 mM]GTP filament end.

**Figure S6.**
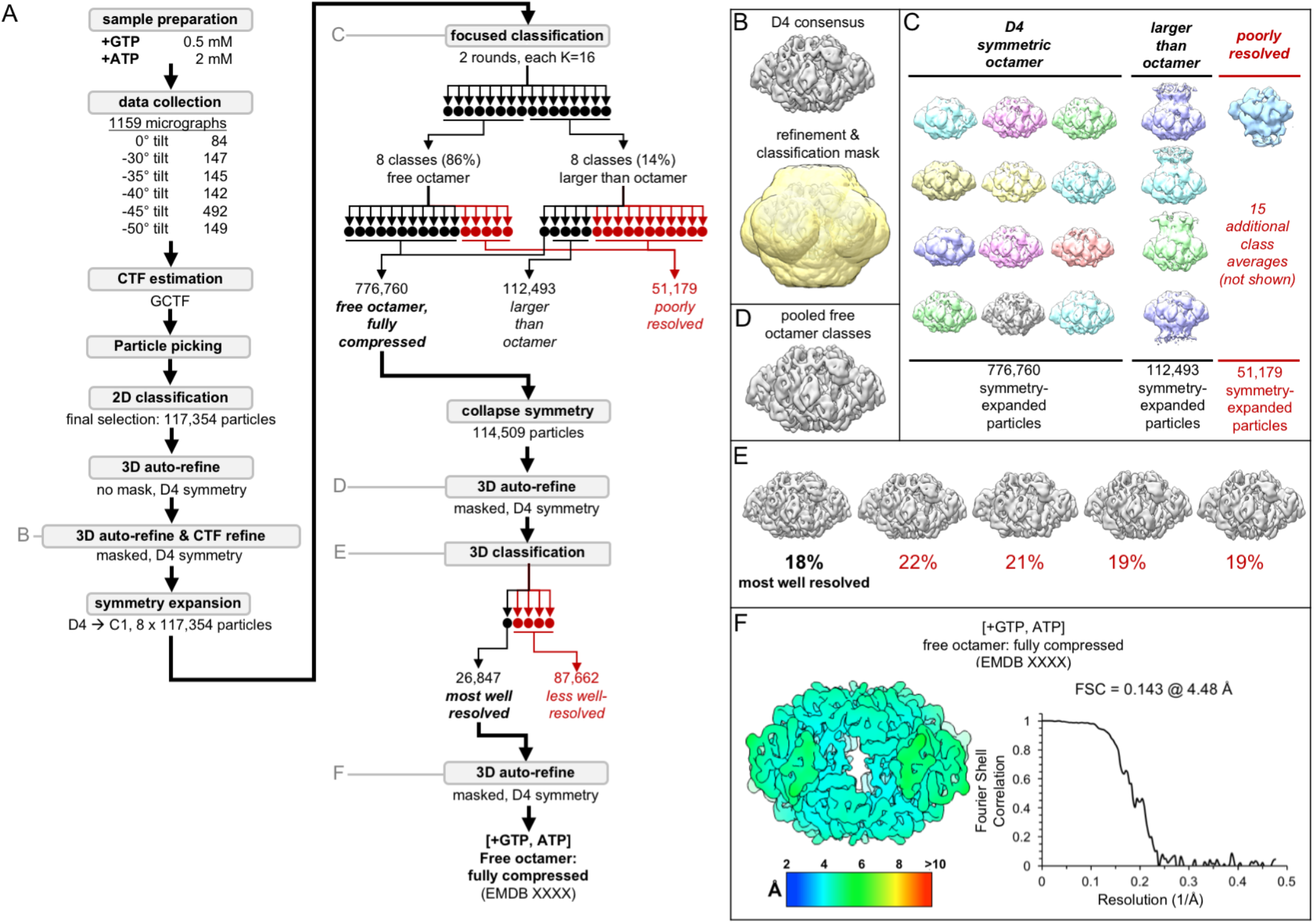
Image processing of the IMPDH2 +ATP, 2 mM GTP cryo-EM dataset. Related to Table 4. A) Flow chart summarizing data processing strategy. B) Masked 3D refinement and all particles from 2D classification/refinement consensus filament assembly interface. Mask also used for all further processing. D4 symmetry enforced during refinement. C) Final class averages from symmetry expanded classification of free octamers. D) Masked refinement from all fully compressed free octamers. E) Final classification of the best-resolved fully compressed free octamers H) Local resolution estimation and FSC curve for the ATP/[2mM]GTP fully compressed free octamer.

**Figure S7.**
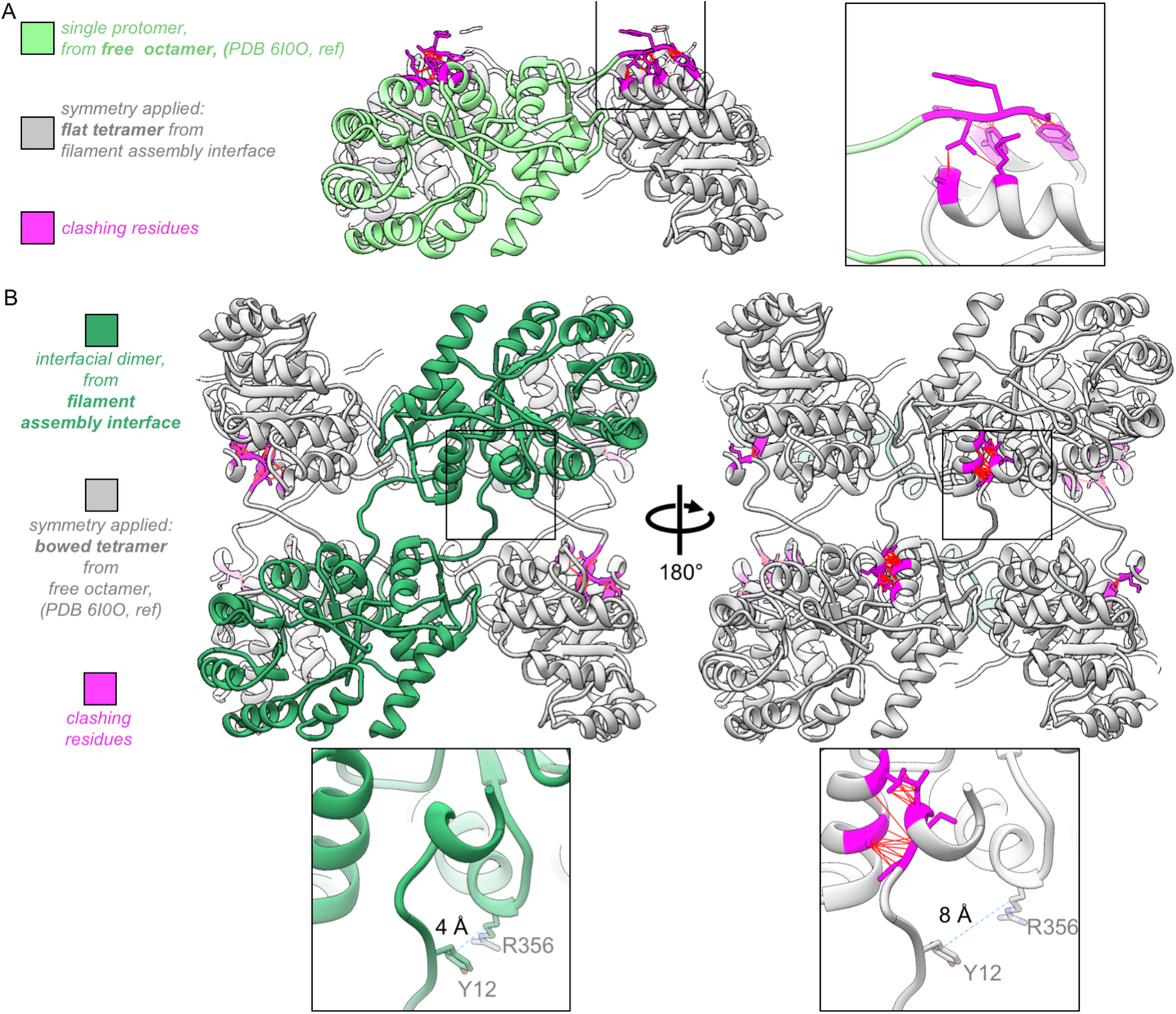
The assembled IMPDH2 filament interface is not compatible with the “bowed” tetramer conformation seen in the unassembled GTP-bound free octamer. Related to Figure 4. A) The protomer from the +GTP crystal structure 6I0O (green), with applied symmetry from the filament assembly interface “flat” tetramer (gray). N-terminus residues that now clash are colored magenta. Inset: closeup of clashing N-terminus. Red lines indicate specific steric clashes. B) Two identical protomers of the filament assembly interface dimer (green) with applied symmetry from the “bowed” tetramer of the +GTP crystal structure 6I0O (gray), with clashing residues colored magenta. Inset: tetramer bowing separates the key residues Y12 and R356 (distances shown are between the gamma carbons of the two residues, indicated by dotted blue line).

**Figure S8.**
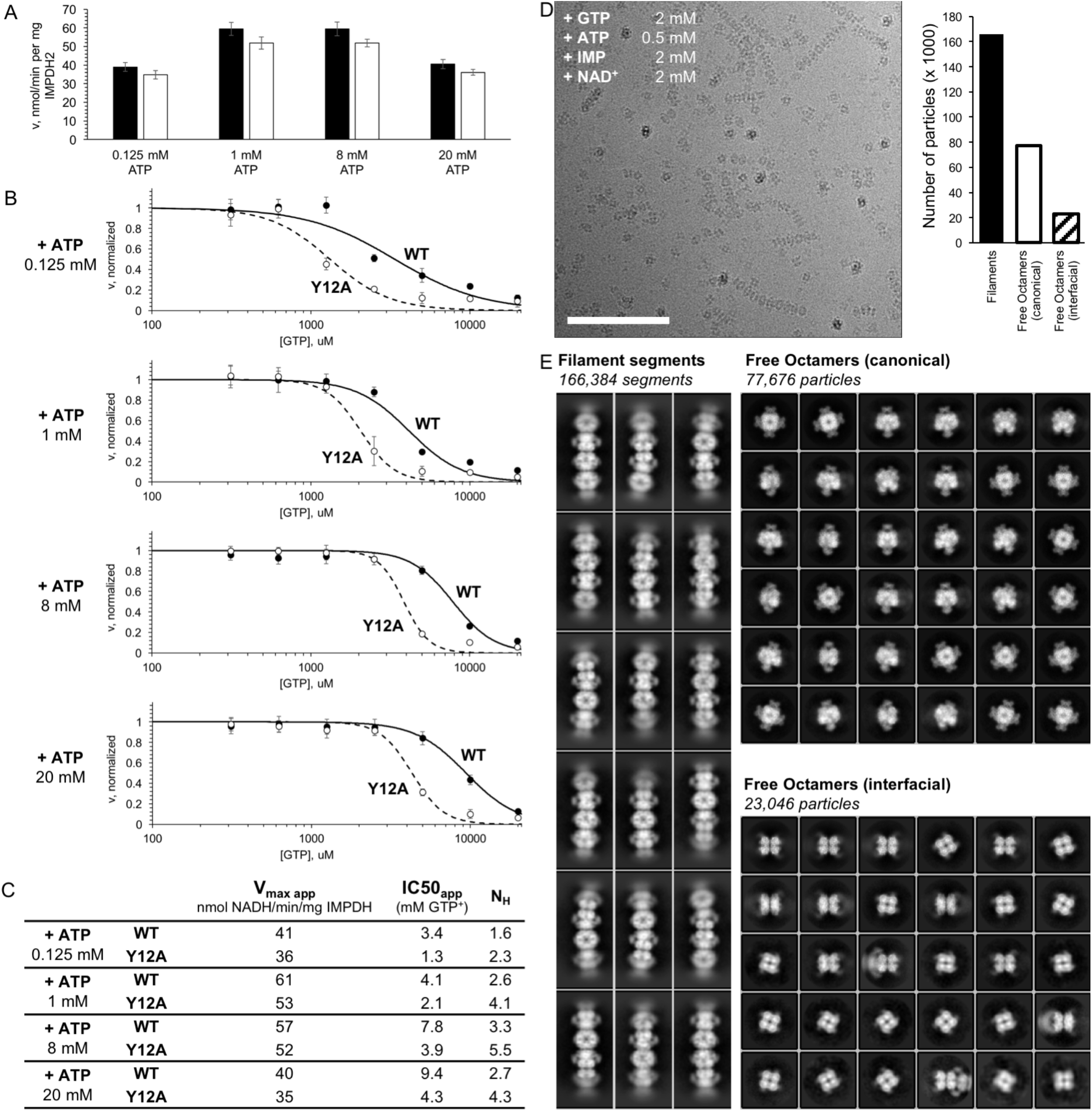
IMPDH2 filaments resist GTP inhibition. Related to Figure 5. A) Initial reaction rates for uninhibited WT enzyme and the filament non-assembly mutant Y12A under various ATP concentrations. High concentrations of ATP likely inhibit by competing with co-substrate NAD^+^. B) Apparent GTP inhibition under saturating substrates (2 mM each IMP and NAD^+^) for a range of ATP concentrations. Under all conditions, WT was more resistant to GTP inhibition than Y12A. C) Estimated Hill equation parameters for data in panel B. D) Cryo-EM of partially inhibited IMPDH2 filaments. Enzyme treated with ATP, IMP, NAD^+^, and 2 mM GTP. Representative micrograph, 2944 total. Scale bar 100 nm. E) Representative 2D class averages of ATP/IMP/NAD+/[2mM]GTP cryo-EM dataset. Three particle types were observed: filament segments, canonical “face-to-face” free octamers, and interfacial “back-to-back” octamers.

**Figure S9.**
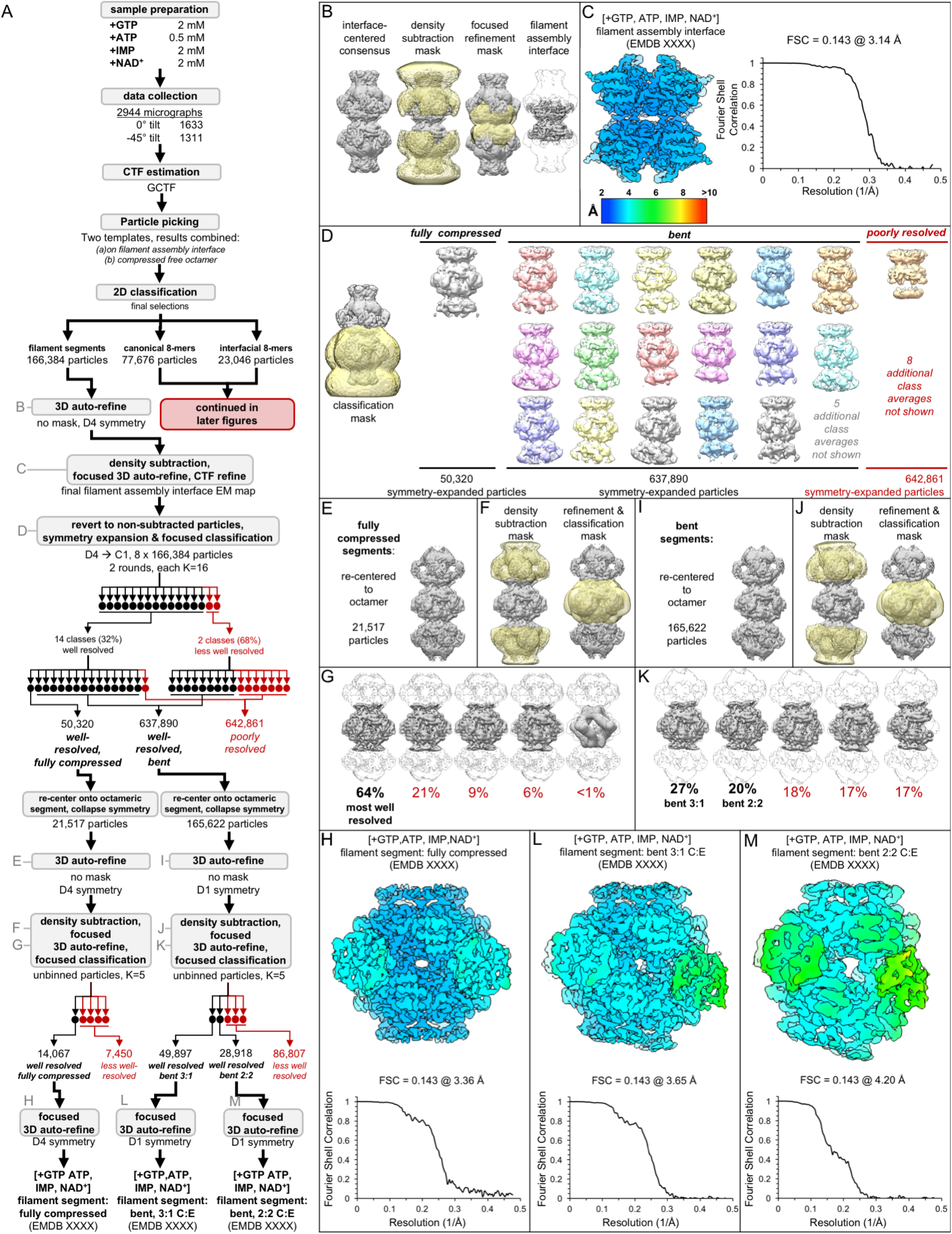
Image processing of the IMPDH2 +ATP, IMP, NAD^+^, 2 mM GTP cryo-EM dataset, part 1: initial processing, and processing of filament segments. Related to Table 6. A) Flow chart summarizing data processing strategy. B) Density subtraction and focused refinement of the consensus filament assembly interface. C) Local resolution estimation and FSC curve (via relion postprocessing) for the ATP/IMP/NAD+/[2mM]GTP consensus filament assembly interface. D) Final class averages from symmetry expanded classification of filament segments. E) Unmasked refinement from all fully compressed segments, pooled and recentered. F) Masks used for continued processing of fully compressed segments. G) Final classification of the best-resolved fully compressed filament segment class H) Local resolution estimation and FSC curve for the ATP/IMP/NAD+/[2mM]GTP fully compressed filament segment I-K) Same as E-G, but for the best-resolved ATP/IMP/NAD+/[2mM]GTP bent filament segment. L-M) Local resolution estimation and FSC curves for the two different ATP/IMP/NAD+/[2mM]GTP bent filament segments.

**Figure S10.**
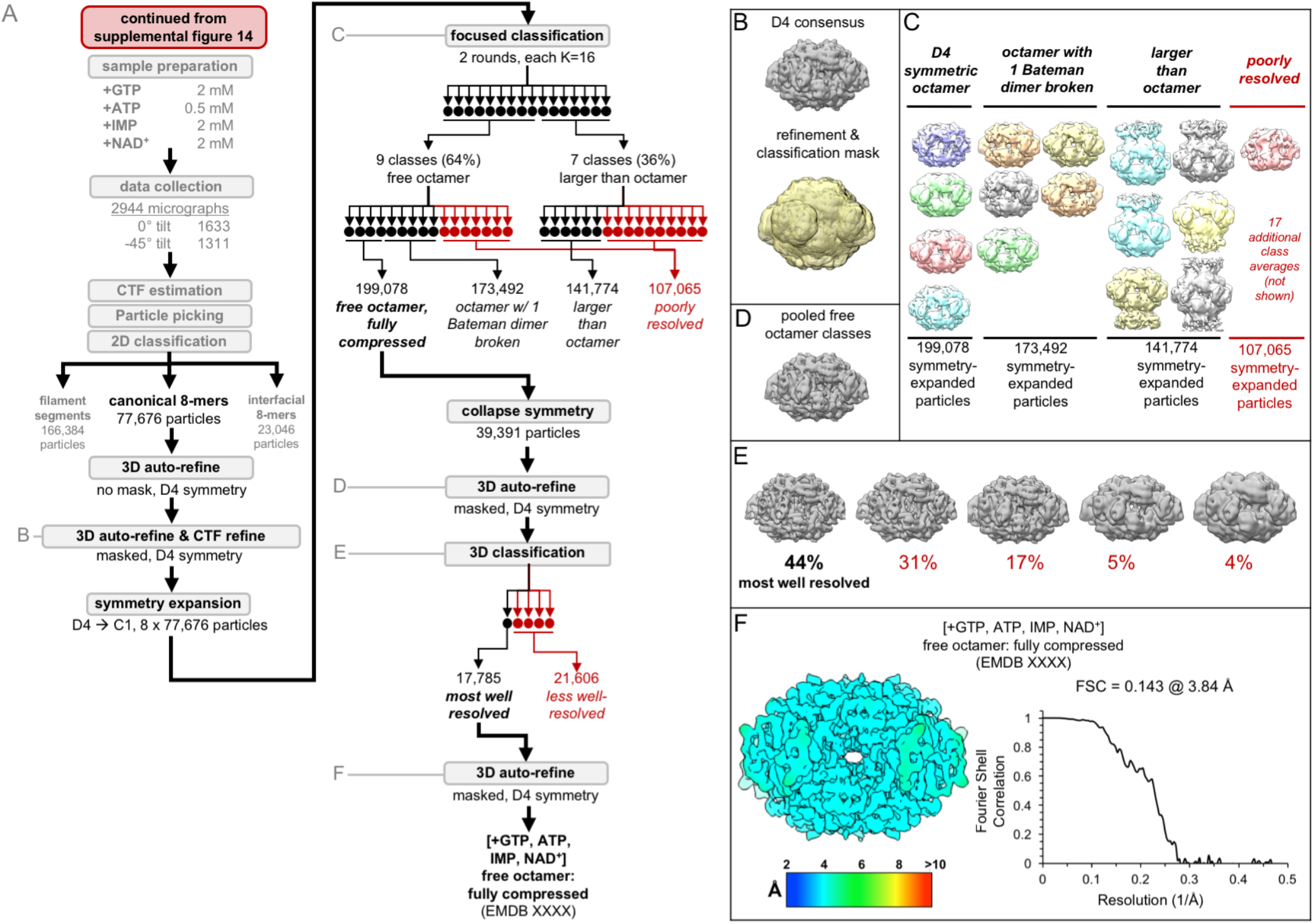
Image processing of the IMPDH2 +ATP, IMP, NAD^+^, 2 mM GTP cryo-EM dataset, part 2: the free canonical octamers. Related to Table 6. A) Flow chart summarizing data processing strategy. B) Masked 3D refinement and all particles from 2D classification/refinement consensus filament assembly interface. Mask also used for all further processing. D4 symmetry enforced during refinement. C) Final class averages from symmetry expanded classification of free octamers. D) Masked refinement from all fully compressed free octamers. E) Final classification of the best-resolved fully compressed free octamers H) Local resolution estimation and FSC curve for the ATP/IMP/NAD^+^/[2mM]GTP fully compressed free canonical octamer.

**Figure S11.**
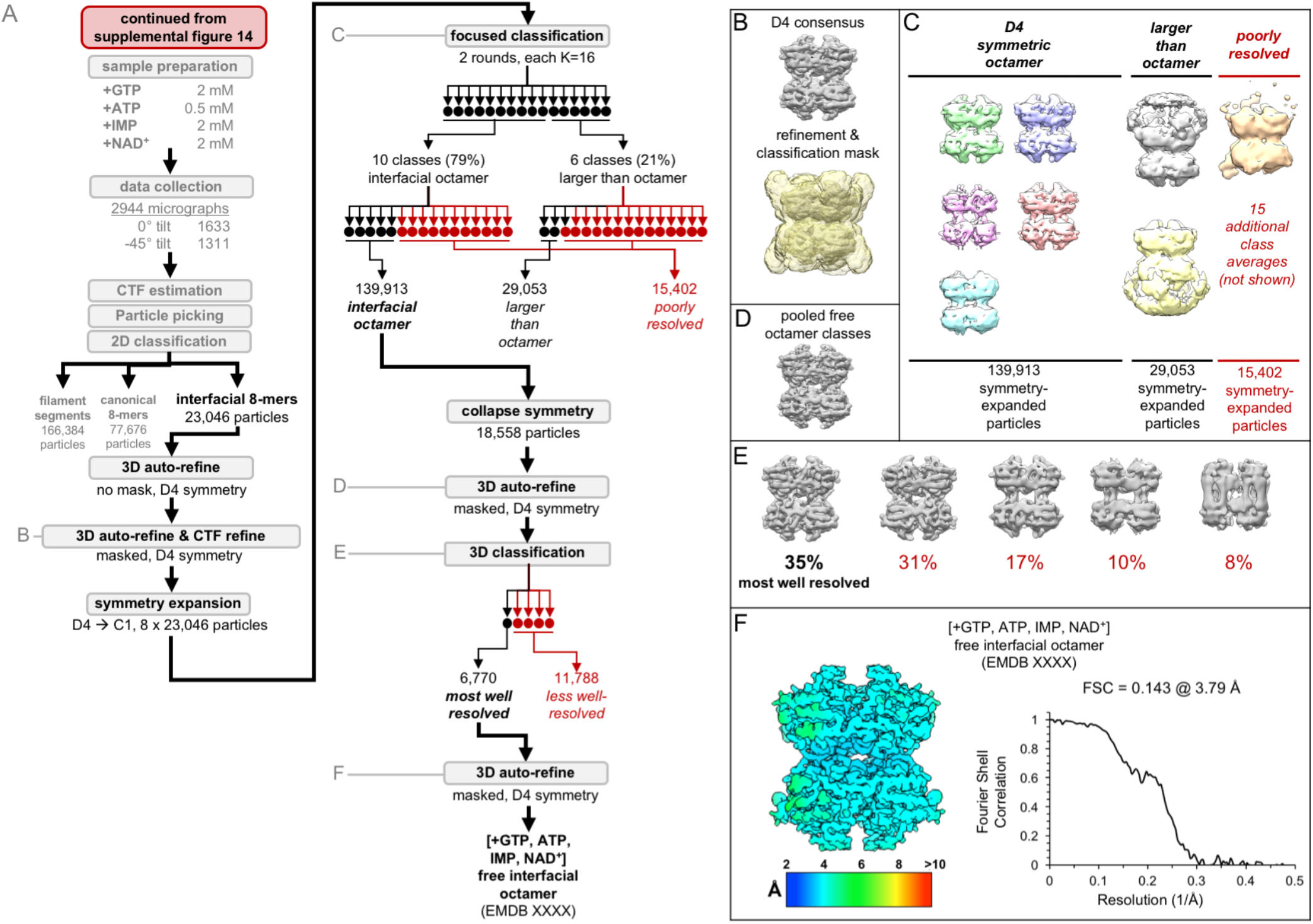
Image processing of the IMPDH2 +ATP, IMP, NAD^+^, 2 mM GTP cryo-EM dataset, part 3: the free interfacial octamers. Related to Table 6. A) Flow chart summarizing data processing strategy. B) Masked 3D refinement and all particles from 2D classification/refinement consensus filament assembly interface. Mask also used for all further processing. D4 symmetry enforced during refinement. C) Final class averages from symmetry expanded classification of free octamers. D) Masked refinement from all fully compressed free octamers. E) Final classification of the best-resolved fully compressed free octamers H) Local resolution estimation and FSC curve for the ATP/IMP/NAD^+^/[2mM]GTP free interfacial octamer.

**Figure S12.**
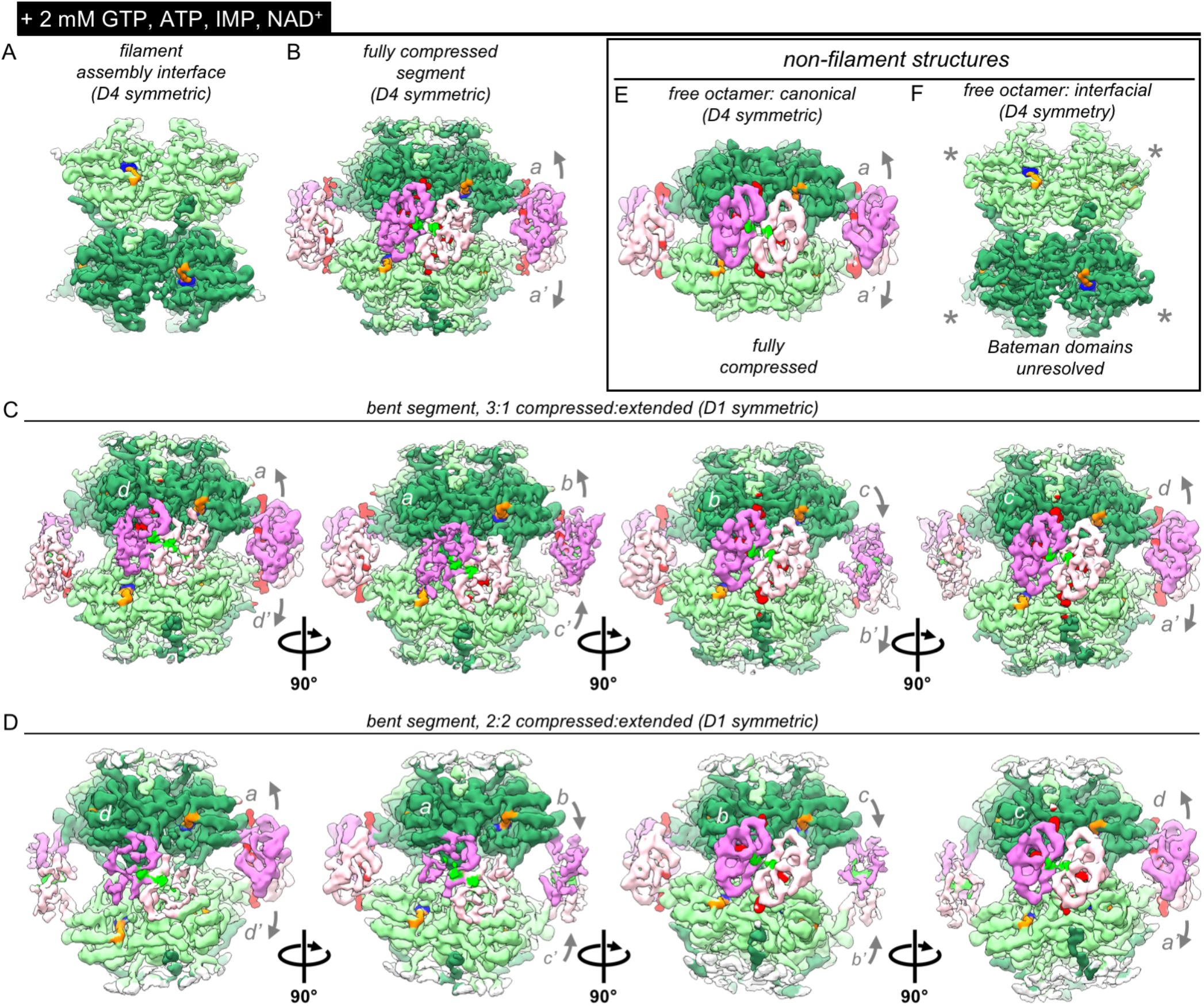
Bateman domains of partially inhibited IMPDH2 filaments are in a mix of compressed and extended states. Related to Figure 5. A) Cryo-EM density of the consensus filament assembly interface from the ATP/IMP/NAD^+^/[2mM]GTP dataset, with density for bound IMP (blue), and NAD^+^ (orange). B) Cryo-EM density of the ATP/IMP/NAD^+^/[2mM]GTP fully compressed filament segment, with (putative) ATP density in Bateman site 1 (bright green) and GTP in sites 2 and 3 (red). C-D) Two D1-symmetric bent filament segments, with a mix of extended and bent protomers. The asymmetric unit of each of these is a tetramer. E-F) Cryo-EM densities of the two different types of D4 symmetric non-filament free octamers resolved from this dataset. The Bateman domains of the free interfacial octamer were unresolved (gray asterisks).

## Methods

### Lead contact and materials availability

Further information and requests for resources and reagents should be directed to and will be fulfilled by the Lead Contact, Justin Kollman.

### Experimental model and subject details

To express recombinant protein constructs, BL21 (DE3) cells acquired from Thermo Scientific were used. For protein purification, the cells were grown in LB broth at 37°C to an optical density of 0.8 and then induced with 1 mM IPTG. After induction, the cells were grown at 30°C for 4 hours. The cells were then harvested using centrifugation.

### Method details

#### Recombinant IMPDH expression and purification

Purified hIMPDH2 was prepared as described previously (Anthony et al. 2017). BL21 (DE3) *E. coli* transformed with a pSMT3-Kan vector expressing N-terminal SMT3/SUMO-tagged hIMPDH2 were cultured in Luria broth at 37°C until reaching an OD_600_ of 0.8 and then induced with 1 mM IPTG for 4 hours at 30°C and pelleted. The remainder of purification was performed at 4°C. Pellets were resuspended in lysis buffer (50 mM KPO4, 300 mM KCl, 20 mM imidazole, 800 mM urea, pH 8) and lysed with an Emulsiflex-05 homogenizer. Lysate was cleared by centrifugation and SUMO-tagged hIMPDH2 chromatagraphically purified with HisTrap FF columns (GE Healthcare Life Sciences) and an Äkta Start chromatography system. After in-column washing with lysis buffer and elution (with 50 mM KPO4, 300 mM KCl, 500 mM imidazole, pH 8), peak fractions were treated with 1 mg ULP 1 protease (Mossessova & Lima 2000) per 100 mg hIMPDH2 for 1 hour, followed by the addition of 1 mM dithiothreitol (DTT) and 800 mM urea. Protein was then concentrated using a 30,000 MWCO Amicon filter and subjected to size-exclusion chromatography using Äkta Pure system and a Superose 6 column pre-equilibrated in filtration buffer (20 mM HEPES, 100 mM KCl, 800 mM urea, 1 mM DTT, pH 8). Peak fractions were flash-frozen in liquid nitrogen and stored at −80°C.

#### IMPDH assembly

Filaments (or, depending on ligand state, free octamers) were prepared by diluting aliquots of purified hIMPDH2 in activity buffer (20 mM HEPES, 100 mM KCl, 1 mM DTT, pH 7) to 2 μM in the presence of varying concentrations of ATP, GTP, IMP, and/or NAD^+^ and incubating for 30 minutes at 20°C.

#### IMPDH activity assays

Protein aliquots were diluted in activity buffer and pre-treated with varying concentrations of ATP, GTP, and IMP for 30 minutes at 20°C, in 96 well UV transparent plates (Corning model 3635). Reactions (100 uL total) were initiated by addition of varying concentrations of NAD^+^. NADH production was measured by optical absorbance (340 nm) in real-time using a Varioskan Lux microplate reader (Thermo Scientific) at 20°C, 1 measurement/min, for 20 minutes; absorbance was correlated with NADH concentration using a standard curve. Specific activity was calculated by linear interpretation of the reaction slope for a 5-minute window beginning 3 minutes after reaction initiation.

#### Negatively stained electron microscopy

Protein preparations were applied to glow-discharged continuous carbon EM grids and negatively stained with 2% uranyl formate. Grids were imaged by transmission electron microscopy using an FEI Tecnai G2 Spirit at 120kV acceleration voltage and a Gatan Ultrascan 4000 CCD using the Leginon software package (Suloway et al. 2009). Micrographs were collected at a nominal 67,000x magnification (pixel size 1.6 Å). GCTF was used for contrast transfer function (CTF) estimation, and Relion for particle picking and 2D classification (Zivanov et al. 2018; Zhang 2016; Scheres 2012).

#### Electron cryo-microscopy sample preparation and data collection

Protein preparations were applied to glow-discharged C-flat holey carbon EM grids (Protochips), blotted, and plunge-frozen in liquid ethane using an Vitrobot plunging apparatus (FEI) at 20°C, 100% relative humidity. High-throughput data collection was performed using an FEI Titan Krios transmission electron microscope operating at 300 kV and equipped with a Gatan image filter (GIF) and post-GIF Gatan K2 Summit direct electron detector using the Leginon software package (Suloway et al. 2009). For the two datasets with non-filament octamers of IMPDH, which exhibit a preferred orientation, it was necessary to collect images with the stage tilted in order to capture a sufficient range of views for 3D reconstruction (Figs. S6A, S9A).

#### Electron cryo-microscopy image processing

Movies were collected in super-resolution mode, then aligned and corrected for beam-induced motion using Motioncor2, with 2X Fourier binning and dose compensation applied during motion correction (Suloway et al. 2009; Zheng et al. 2017). CTF was estimated using GCTF (Zhang 2016). Relion 3.0 was used for all subsequent image processing (Zhang 2016; Zivanov et al. 2018). Although multiple datasets of hIMPDH2 under the different ligand states were collected, each dataset was individually processed using approximately the same overall pipeline (Figs. S1E-G), with some variations from dataset to dataset (Figs. S2, S5-6, S9-11). First, for each dataset, autopicking templates and initial 3D references maps were prepared by manually picking and extracting boxed particles from a small subset of micrographs, and classifying/refining in 2D and 3D. For these initial 3D refinements, a featureless, soft-edged cylinder was used as a refinement template of filaments, and a previously published cryo-em map (EMDB-8692) was used as template for non-filament octamers (Anthony et al. 2017). Because IMPDH filament segments possess D4 point-group symmetry, two different locations along filaments may be used as symmetry origins: the centers of canonical octamer segments, or the centers of the assembly interface between segments. For the filament datasets, we prepared and used auto-picking templates centered on the filament assembly interface. For the datasets containing non-filament octamers of hIMPDH2, auto-picking templates centered on these non-filament particles were also included. Due to the flexibility of hIMPDH2 filaments, helical segments were processed as single particles, and at no point was helical symmetry applied during image processing. After template-based autopicking of each complete dataset, picked particles were boxed and extracted from micrographs, and subjected to hierarchical 2D classification to select the best-resolved classes. These selected particles were then auto-refined in 3D as a single class with symmetry applied (D4 for filament segments and free octamers, C4 for filament ends). Exploratory image processing of the assembly interface-centered filament reconstructions made it apparent that the eight catalytic domains surrounding this interface appeared conformationally homogenous, while the Bateman domains and neighboring octamers appeared conformationally varied. Additionally, due to the flexibility of the filaments, and the tendency of filament ends to adhere to the air-water interface, many filaments were tilted out of plane, with neighboring segments overlapping in projection.

To improve resolution, partial signal subtraction was performed at this stage, using a mask that left only the central eight catalytic domains of the filament assembly interface, subtracting the poorly resolved Bateman domains and neighboring segments, which served to improve resolution after subsequent auto-refinement. Per-particle defocus and per-micrograph astigmatism were then optimized using CTF refinement, which improved resolution further. The resulting consensus refinements of the filament assembly interfaces were well-resolved, however data on Bateman domain conformation was missing, with these regions very poorly resolved when subtracted regions were restored to the reconstructions by reversion to original non-subtracted particles (data not shown). To resolve the different Bateman domain conformations, we applied particle symmetry expansion (D4 to C1) and classified particles without additional alignment. Because at this stage the reconstructions were centered on the filament assembly interface, each boxed “particle” contained elements of two different neighboring octamers. The potential conformational space was reduced by applying a mask enclosing only one of these two octamers. By hierarchical focused classification of the off-origin octamers we were able to classify multiple conformations of the octameric filament segments, as well as incomplete segments and filament ends. Symmetry expansion was also applied to the non-filament octamer datasets, with a mask including the entire particle, which allowed classification of the most symmetric and well-resolved classes. To further improve resolution of the varying symmetry-expanded segment classes, the reconstruction symmetry origins were moved from the filament assembly interface to the canonical octamers by re-extraction with re-centering.For each class, symmetry was then collapsed by removing redundant overlapping particles, Euler angles reset to zero. After auto-refining once again, we observed that the most well-resolved octameric segments from the asymmetric symmetry-expanded classifications exhibited some apparent symmetry, with fully extended or fully compressed octamers appearing D4 symmetric, and some bent classes apparently D1 symmetric. We therefore applied these symmetries during subsequent refinement and classification of these new octamer-centered classes. As before, signal subtraction of neighboring filament segments improved resolution considerably. Additional rounds of CTF refinement and 3D classification identified the best-resolved particles from each of these conformational classes. Final overall resolution (according to the FSC=0.143 criterion), as well as local resolution, was assessed using Relion postprocessing.

#### Model building and refinement

As initial templates for model building, two hybrid models (representing hIMPDH2 in either an extended or compressed state) were prepared by combining elements from existing crystal structures. For both templates, the catalytic domain and substrate poses (residues 18-107, & 245-514) were taken from a crystal structure of an inhibitor-bound hIMPDH2 (PDB 1nf7), and the Bateman domains and ligand poses (residues 108-244) were based on fungal (*A. gossypii)* IMPDH crystallized in either the extended or compressed states (PDB 5mcp and 5tc3, respectively), and SWISS-MODEL homology modeling (Sintchak et al. 1996; Waterhouse et al. 2018; Buey et al. 2017). The N-terminus (residues 1-17) were modelled by hand. In all maps, a single active site loop (residues 421 to 436) was unresolved, and these residues were not modelled. After rigid-body fitting of templates into the cryo-EM densities using UCSF Chimera, repeated cycles of manual fitting with Coot, automated fitting with phenix.real_space_refine (employing rigid-body refinement, NCS constraints, gradient-driven minimization and simulated annealing) and local B-factor sharpening of cryo-EM data via LocScale were used for final atomic model refinement and local sharpening of cryo-EM maps (Pettersen et al. 2004; Emsley et al. 2010; Adams et al. 2012; Jakobi et al. 2017). Final models were evaluated with MOLPROBITY and EMRinger (Chen et al. 2010; Barad et al. 2015). Data collection parameters and refinement statistics are summarized in Tables 1-6. Figures were prepared with UCSF Chimera (Pettersen et al. 2004).

### Quantification and statistical analysis

Protein concentrations were assayed with a NanoDrop spectrophotometer (Thermo Scientific). NADH concentrations were assayed with a Varioskan Lux microplate reader (Thermo Scientific). Per-residue backbone RMSD values were calculated using superposed models in UCSF Chimera. All statistical validation performed on the deposited maps/models was done using Relion, PHENIX, MolProbity, and EMringer.

### Data and code availability

The cryo-EM maps described here have been deposited in the Electron Microscopy Data Bank with accession numbers 20687, 20688, 20690, 20691, 20701, 20704, 20705, 20706, 20707, 20709, 20716, 20718, 20720, 20722, 20723, 20725, 20742, 20741, and 20743. The refined atomic coordinates have been deposited in the Protein Data Bank with accession numbers 6U8E, 6U8N, 6U8R, 6U8S, 6U9O, 6UA2, 6UA4, 6UA5, 6UAJ, 6UC2, 6UDP, 6UDO, and 6UDQ.

## Supplemental Items

**Video S1. Comparison between the “flat” tetramer of assembled filaments and the “bowed” tetramer of free octamers.** Related to Figure 4. Morph comparison between catalytic tetramers of the GTP/ATP/IMP filament assembly interface and the GTP/ATP free octamer. For visualization purposes, we have depicted the complete N-terminus in both conformations, however in the free octamer model, some residues were unresolved (gray). For both models, opposing tetramers, Bateman domains, and active site loops have been hidden from view.

